# Mammalian RNAi represses pericentromeric lncRNAs to maintain genome stability

**DOI:** 10.1101/2024.05.09.593425

**Authors:** Rafael Sandoval, Corinne N. Dilsavor, Nadia R. Grishanina, Vandan Patel, Jesse R. Zamudio

## Abstract

Mammalian pericentromeric tandem repeats produce long noncoding RNAs (lncRNAs) that are dysregulated in cancer and linked to genomic instability. Identifying the basic molecular characteristics of these lncRNAs and their regulation is important to understanding their biological function. Here, we determine that the Argonaute (Ago) proteins of the RNA interference (RNAi) pathway directly and uniformly repress bidirectional pericentromeric lncRNAs in a Dicer-dependent manner in mouse embryonic and adult stem cells. Ago-dependent and Dicer-dependent autoregulatory small RNAs were identified within pericentromeric lncRNA degradation intermediates. We develop an RNase H cleavage assay to determine the relative proportions and lengths of the pericentromeric lncRNA targets. We find that 5’-phosphate and non-polyadenylated bidirectional pericentromeric lncRNAs are expressed at similar proportions. These lncRNAs can span up to 9 repeats, with transcription from the reverse strand template yielding the longer products. Using pericentromeric repeat RNA reporters, we determine that Ago represses pericentromeric lncRNAs after S phase transcription. Upon loss of Ago, pericentromeric lncRNA dysregulation results in delayed cell cycle progression, a defective mitotic spindle assembly checkpoint (SAC) and genomic instability. These results show that an evolutionarily conserved Ago activity at pericentromeres contributes to mammalian genome stability.

## Introduction

Pericentromeres are chromosomal regions containing large tandem repeat arrays that serve an important role in cell division (Müller and Almouzni 2017). These repeats are found within constitutive heterochromatin and act in sister chromatid cohesion. This cohesion activity at pericentromeres contributes to genome stability by preventing erroneous chromosome separation before the metaphase to anaphase transition in mitosis. The transition to anaphase is a critical stage in the eukaryotic cell cycle and is regulated by the mitotic SAC (Lara-Gonzalez et al. 2012). Defects in either pericentromeric heterochromatin (Peters et al. 2001) or the SAC (Li and Murray 1991) result in genomic instability that can contribute to cancer.

RNAs transcribed from the pericentromeric repeats are also implicated in genome stability. Pericentromeric RNAs promote progression through early embryonic stem cell divisions (Probst et al. 2010) and may act directly in local heterochromatin formation (Smurova and Wulf 2018). These RNAs are overexpressed in human epithelial cancers (Ting et al. 2011), and their dysregulation advances tumor formation in murine cancer models (Zhu et al. 2011) (Zhu et al. 2018). Mechanistically, the forced overexpression of pericentromeric RNA in these models caused dysregulation of SAC activity and increased genomic instability. Although linked to mammalian development and disease, the basic molecular properties of the pericentromeric RNAs remain unclear because of the challenges in characterizing RNA from large tandem repeats.

In fission yeast and multiple other non-mammalian model systems, RNAi functions at pericentromeres through autoregulatory small RNAs and contributes to genome stability (Volpe et al. 2002; Pek and Kai 2011; Claycomb et al. 2009; Mochizuki and Gorovsky 2005; Durand-Dubief and Bastin 2003). RNAi is a gene regulatory pathway in which effector Ago proteins are guided by small RNAs to regulate RNA targets with complementary sequences (Ipsaro and Joshua-Tor 2015). In *S. pombe*, bidirectional transcripts from the pericentromeres are substrates for Dicer nuclease processing into autoregulatory small RNAs that guide Ago for local heterochromatin formation (Martienssen and Moazed 2015). Whether RNAi directly functions at pericentromeres in mammals remains controversial (Gutbrod et al. 2022), and a detailed characterization of Ago in mammalian genome stability through cell division is lacking.

Here, we used conditional Ago-expressing mouse embryonic stem cells (mESCs) (Zamudio et al. 2014; Bosson et al. 2014; Kelly et al. 2019b, 2019a; JnBaptiste et al. 2017) to address RNAi regulation at pericentromeres. We found that pericentromeric lncRNAs are direct Dicer-dependent targets of Ago repression. The pericentromeric lncRNA targets are bidirectional, proportionally transcribed, chromatin-associated, heterogeneous in repeat number, and contain termini that indicate distinct RNA processing events. We determined that Dicer- and Ago-dependent small RNAs are generated from degradation intermediates derived from the bidirectional pericentromeric lncRNAs. Upon Ago or Dicer loss, bidirectional pericentromeric lncRNAs are uniformly upregulated. Ago repression occurs after S phase transcription and failure to suppress these lncRNAs results in defective cell cycle progression and abnormal mitotic chromosome segregation. This study uncovers the RNA basis for Ago gene repression at mammalian pericentromeres and defines its role in genome stability.

## Results

### The RNAi pathway uniformly represses bidirectional pericentromeric lncRNAs

Similar to humans, mouse pericentromeres are composed of large tandem DNA repeats that function as important structural elements for the control of mitotic chromosome segregation (McKinley and Cheeseman 2016). The mouse pericentromere major satellite repeat (MSR) is 234 bp long and can be found in over 10,000 tandem repeats on each chromosome (Vissel and Choo 1989). Within the MSR, four major satellite subrepeats (MSS) show high sequence similarity and contain promoters for bidirectional transcription (Fig. 1A) (Bulut-Karslioglu et al. 2012). In *S. pombe*, long pericentromeric transcripts appear to be unstable scaffolds acting in the local recruitment of RNAi factors. The current approaches to detect mammalian pericentromeric lncRNAs do not permit a clear assessment of the RNA products and their potential regulation. This leaves an important open question in cellular biology regarding whether bidirectional transcription at pericentromeres triggers RNAi regulation in mammalian cells.

**Figure 1.**
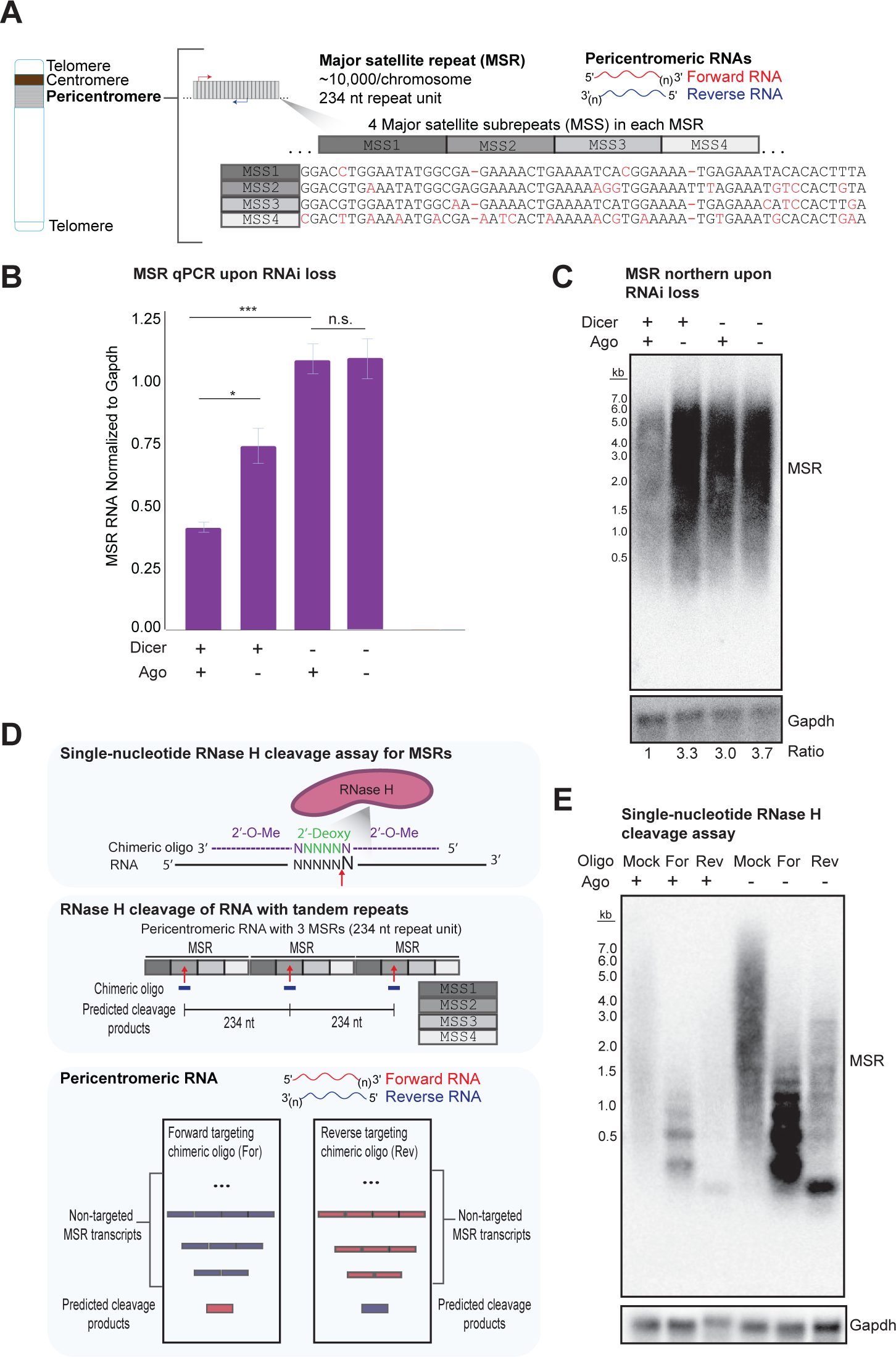
The RNAi pathway uniformly represses bidirectional pericentromeric lncRNAs. (A) Diagram of the mouse pericentromere. Alignment of the four major satellite subrepeats (MSS) with nucleotide differences indicated in red. Bidirectional transcription is indicated by arrows and transcripts colored in red (forward) or blue (reverse). (n) indicates the uncertainty of repeat numbers in pericentromeric lncRNAs. (B) RT-qPCR in Dicer Het or KO cells treated for Ago2 (Dox) or No Ago (No Dox) expression. (*: p value < 0.05; ***: p value < 0.001; n.s.: nonsignificant; p value determined by two-sided t-test of biological triplicates) (C) Northern of Dicer Het or KO cells treated for Ago2 (Dox) or No Ago (No Dox) expression. Top: Random hexamer probe for MSRs; Bottom: Random hexamer probe for Gapdh. Ratio of MSR to Gapdh indicated below. (D) Diagram of RNase H cleavage assay for specific targeting and detection of pericentromeric lncRNAs. (E) Northern of total RNA from cells treated for Ago2 (Dox) or No Ago (No Dox) expression. Pericentromeric RNA was targeted for cleavage by RNase H guided by either a forward (For) or reverse (Rev) MSR targeting chimeric oligo or mock (no chimeric oligo) treatments. Top: Random hexamer probe for MSR; Bottom: Random hexamer probe for Gapdh.

To characterize RNAi regulation of pericentromeric lncRNAs, we used conditional Ago-expressing mESCs that express a FLAG and HA epitope-tagged Ago2 gene in an Ago1-4 knockout background, allowing complete characterization of Ago activity (Zamudio et al. 2014). We engineered Dicer knockouts (KO) in this conditional Ago-expressing cell line to determine Dicer-dependent Ago regulation. We first measured MSR RNA in RNAi mutants by qPCR, which did not differentiate RNA strands. We found that Ago depletion in Dicer-expressing cells increased MSR RNA detection (Fig. 1B). In Dicer-deficient cells, MSR detection was also increased supporting Ago- and Dicer-dependent regulation of pericentromeric RNAs. Since increased qPCR detection does not differentiate between greater RNA abundance and more repeat numbers within transcripts, we set out to visualize the target RNA populations. As previously reported (Lu and Gilbert 2007), we observed RNA signal from MSRs showing size heterogeneity spanning ∼0.5 kb-7 kb in northern blots probed for MSR forward and reverse strands (Fig. 1C). We detected a ∼3-4 fold increase of the pericentromeric lncRNA signal upon Ago or Dicer depletion relative to the loading control, and no other differences in the distribution of the RNA signal. These results indicate that RNAi uniformly suppresses the abundance of a heterogeneous population of pericentromeric lncRNAs.

To further characterize the pericentromeric lncRNAs, we assayed RNA processing events. First, we measured protection from the 5’-phosphate-dependent 5’→3’ exonuclease Xrn1. We found that pericentromeric lncRNAs were susceptible to Xrn1 degradation, in contrast to m^7^G-capped Gapdh mRNA (Supplemental Fig. S1A). Next, we performed polyA RNA pulldowns and detected pericentromeric lncRNAs in the unbound fraction, in contrast to the mRNA control (Supplemental Fig. S1B). These results indicate that pericentromeric lncRNAs contain a 5’ phosphate and lack a polyA tail. Consistent with this characterization, we detected higher read counts for bidirectional pericentromeric lncRNAs in total RNAseq libraries (data not shown) and determined upregulation of RNA from both forward and reverse strands upon Ago or Dicer depletion (Supplemental Fig. S1C).

To further define the pericentromeric lncRNAs targeted by RNAi, we developed an RNase H single-nucleotide cleavage assay (Lapham and Crothers 2000). This RNase H approach allows the prediction of cleavage products that validate the specificity of pericentromeric lncRNA detection and permits strand-specific quantification (Fig. 1D). The RNase H cleavage of transcripts containing tandem repeats should collapse signal near the repeat length, while non-targeted RNA should be unaltered. We designed RNase H chimeric oligonucleotides to cleave forward or reverse pericentromeric lncRNAs at the region of highest sequence divergence between subrepeats. Using the chimeric oligonucleotide that targets forward transcripts, we detected a shift in RNA signal from the untreated ∼0.5 kb-7 kb smear to four distinct bands < 1.5 kb (Figure 1E). This RNase H cleavage pattern validated the detection of RNA and not contaminant DNA, while the smallest band at ∼234 nt confirmed northern probe specificity for pericentromeric transcripts. Interestingly, the larger bands migrated distinctly from the previously observed smears and were instead separated by the ∼234 nt repeat unit size. This result reveals the sizes and relative proportion of reverse pericentromeric lncRNAs. Specifically, reverse pericentromeric lncRNAs are generated or processed to contain 1-4 MSRs and are expressed at similar levels to forward pericentromeric lncRNAs. The classification of these RNA species was confirmed with directional oligonucleotide probes (Supplemental Fig. S1D).

Using the chimeric oligonucleotide that targets reverse transcripts, we revealed the sizes of the forward pericentromeric lncRNAs, determining that they exist as longer species with 2-9 repeats. Upon depletion of Ago, both forward and reverse transcripts were uniformly upregulated with no apparent change in RNA length or relative forward and reverse proportions. The products of the RNase H assay were confirmed on higher resolution northern blots and by fragment sequencing (Supplemental Fig. S1E,F). This RNase H assay provides clarity of bidirectional pericentromeric lncRNA size, orientation, and proportions and demonstrates their uniform regulation by RNAi.

### Ago- and Dicer-dependent autoregulatory small RNAs from pericentromeric lncRNAs

To search for evidence of direct Ago regulation of the pericentromeric lncRNAs, we aimed to define autoregulatory small RNA guides, as characterized in other model systems. Small RNA sequencing libraries are a mixture of Ago-bound small RNA guides and fragments from RNA degradation. It is analytically challenging to distinguish autoregulatory guides from other degradation fragments generated during the decay of primary RNA transcripts. To overcome this challenge, we identified Ago guides based on Dicer-dependent processing and Ago-dependent stability as previously reported for known guides such as microRNA (miRNA) (Fig. 2A) (Zamudio et al. 2014). We devised an approach based on grouped size ranges, including those expected to contain RNAi guides (20-25mers) and other size ranges expected to be mostly degradation fragments. First, we determined that Dicer-expressing mESCs have higher amounts of forward and reverse 20-25mer fragments from MSRs relative to other fragment sizes (Fig. 2B). The 20-25mers were reduced in Dicer-deficient cells indicating a subpopulation that is Dicer-dependent. We further characterized the small RNA profile differences in Dicer- and Ago-deficient mESCs. We found that 20-25mers from both forward and reverse pericentromeric repeats were less abundant in Dicer-deficient and Ago-deficient cells compared to cells expressing these proteins (Fig. 2C). As expected, Dicer or Ago loss did not shift the degradation profile of H/ACA small nucleolar RNAs (snoRNAs), but largely decreased the proportions of miRNAs. These results indicate specific small RNA changes in pericentromeric RNAs that are consistent with the generation of autoregulatory guides for Ago.

**Figure 2.**
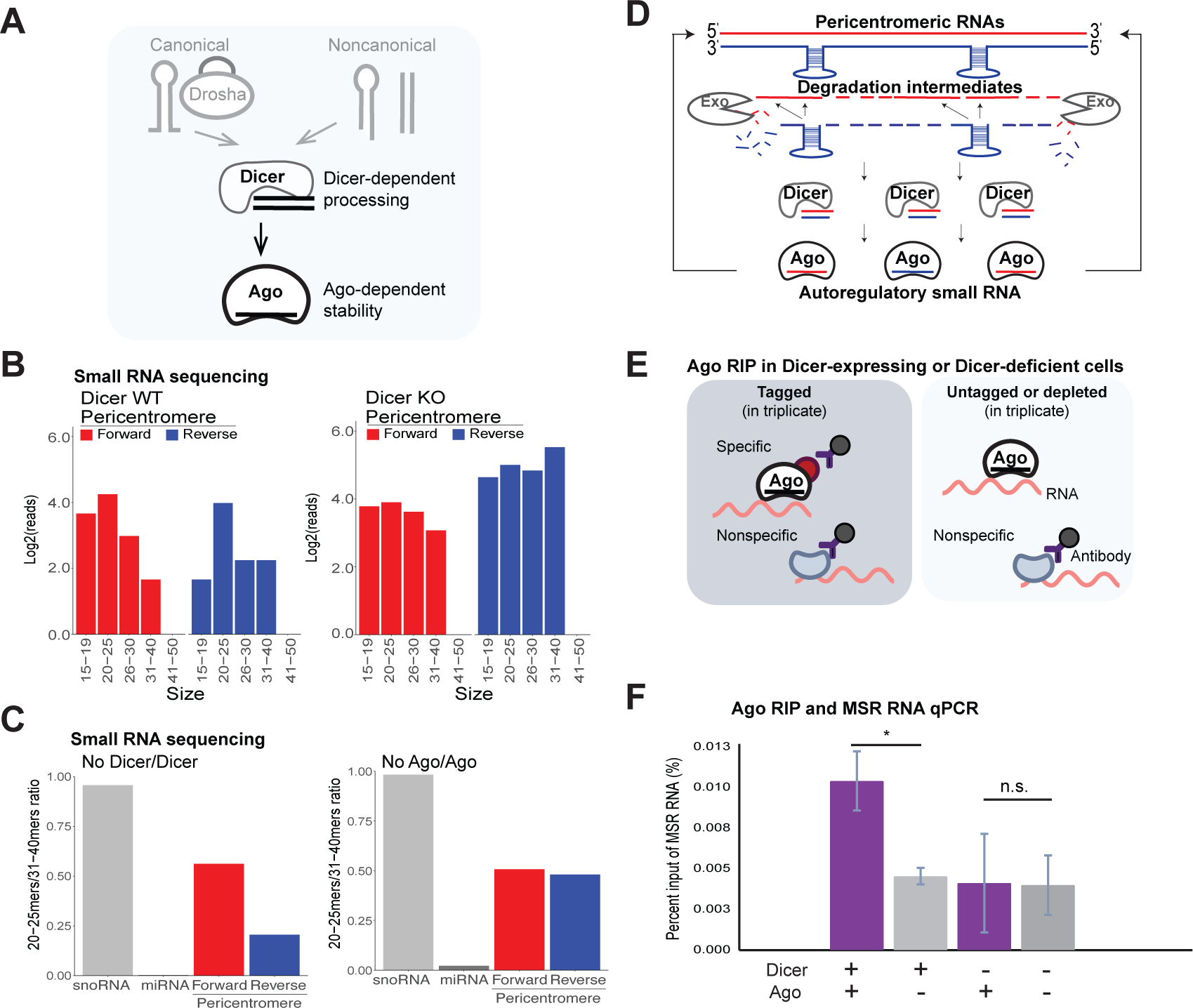
Ago- and Dicer-dependent autoregulatory small RNAs from pericentromeric lncRNAs. (A) Diagram of RNAi pathway indicating detection criteria for autoregulatory small RNAs: Dicer- and Ago-dependent classification. (B) Barplot of MSR reads from small RNA sequencing libraries grouped by indicated size ranges in Dicer WT (left) or Dicer KO (right) mESCs. On each plot forward (red) and reverse (blue) counts are indicated. (C) The ratio of 20-25mers to 31-40mers was calculated for H/ACA box snoRNAs, miRNAs, and the forward (red) and reverse (blue) MSR. Left: Dicer KO/Dicer WT; Right: No Ago/Ago2 expression conditions. (D) Model for autoregulatory small RNAs produced from pericentromeric lncRNA degradation intermediates. (E) Diagram of Ago RIP experiments with negative controls for antibody specificity and Dicer-dependence. (F) RIP-qPCR for Dicer Het or Dicer KO cells treated for Ago2 (Dox) or No Ago (No Dox) expression. The qPCR conditions optimized for this assay detect a 151 bp region that spans two repeat units and uses a short extension time to generate a single amplicon. (*: p value < 0.05; n.s.: nonsignificant; p value determined by two-sided t-test of biological triplicates)

To gain further insight into these autoregulatory guides, we mapped the 5’ start positions of small RNAs along the MSR (Supplemental Fig. S2A). We observed four MSR forward clusters and three reverse clusters skewed toward the 3’ end of MSS regions. The distribution of the size proportions at each cluster indicates local degradation by exonucleases (Supplemental Fig. S2B). We hypothesized that the detection of these clusters compared to flanking sequences likely relates to structural elements delaying degradation, such as hairpins or short double-stranded RNAs (dsRNAs). Interestingly, secondary structure predictions did not indicate hairpins of sufficient length to generate 20-25mers (Supplemental Fig. S2C,D). However, the reverse RNA structure prediction indicated short hairpins in G-U rich regions that would not form such hairpins in the C-A complementary sequences. These reverse hairpins may delay RNA degradation, and these degradation intermediates anneal to multiple MSS positions within forward sequences to produce dsRNA for Dicer processing (Fig. 1D; Supplemental Fig. S2F).

Because this model allowed the prediction of Ago-bound small RNA positions along the pericentromeric repeat, we focused on higher resolution profiling of Ago-dependent small RNAs. We found that 20-25mers decreased in Ago-deficient cells within forward and reverse clusters, indicating Ago-bound small RNAs from the predicted dsRNA positions (Supplemental Fig. S2G). To confirm this, we examined the pericentromeric small RNAs in Ago2 small RNA immunoprecipitation (RIP) experiments. We determined a skew toward MSR 20-25mers in epitope-tagged Ago2 RIPs compared to untagged Ago2 negative controls, and within the observed RNA clusters (Supplemental Fig. S2H,I). Overall, the small RNA profiles fit a model of Ago-bound small RNAs generated by Dicer processing of degradation intermediates from bidirectional transcription.

Since the small RNA characterization indicated the production of autoregulatory guides by Dicer, we predicted Dicer-dependent association of Ago with pericentromeric lncRNAs. Consistent with this, we found that Ago binding to pericentromeric lncRNAs is Dicer-dependent as assayed by Ago RIP followed by RT-qPCR in Dicer-expressing or Dicer-deficient cells (Fig. 2E,F). Similar pericentromeric RNA results were determined in small RNA sequencing and Ago RIPs in adult mouse mesenchymal stem cells (mMSCs) (JnBaptiste et al. 2017) (Supplemental Fig. S3). Together, these results suggest that Ago uses autoregulatory guides to repress the longer pericentromeric lncRNAs in both embryonic and adult stem cells.

### MSRs impart Ago-mediated regulation on an exogenous reporter

We next confirmed that transcription from MSRs can drive RNAi regulation while being detected within constitutive heterochromatin. We reasoned that the ability to introduce MSRs into the genome that cause heterochromatin formation, transcription and RNAi regulation would support regions targeted for all three activities. To this end, we developed a major satellite repeat mCherry single-cell RNA (MSR2) reporter. We designed the MSR2 reporter with an internal ribosome entry site (IRES)-mCherry-pA downstream of nine MSRs (Fig. 3A). The MSR2 reporter was stably integrated into conditional Ago-expressing mESCs.

**Figure 3.**
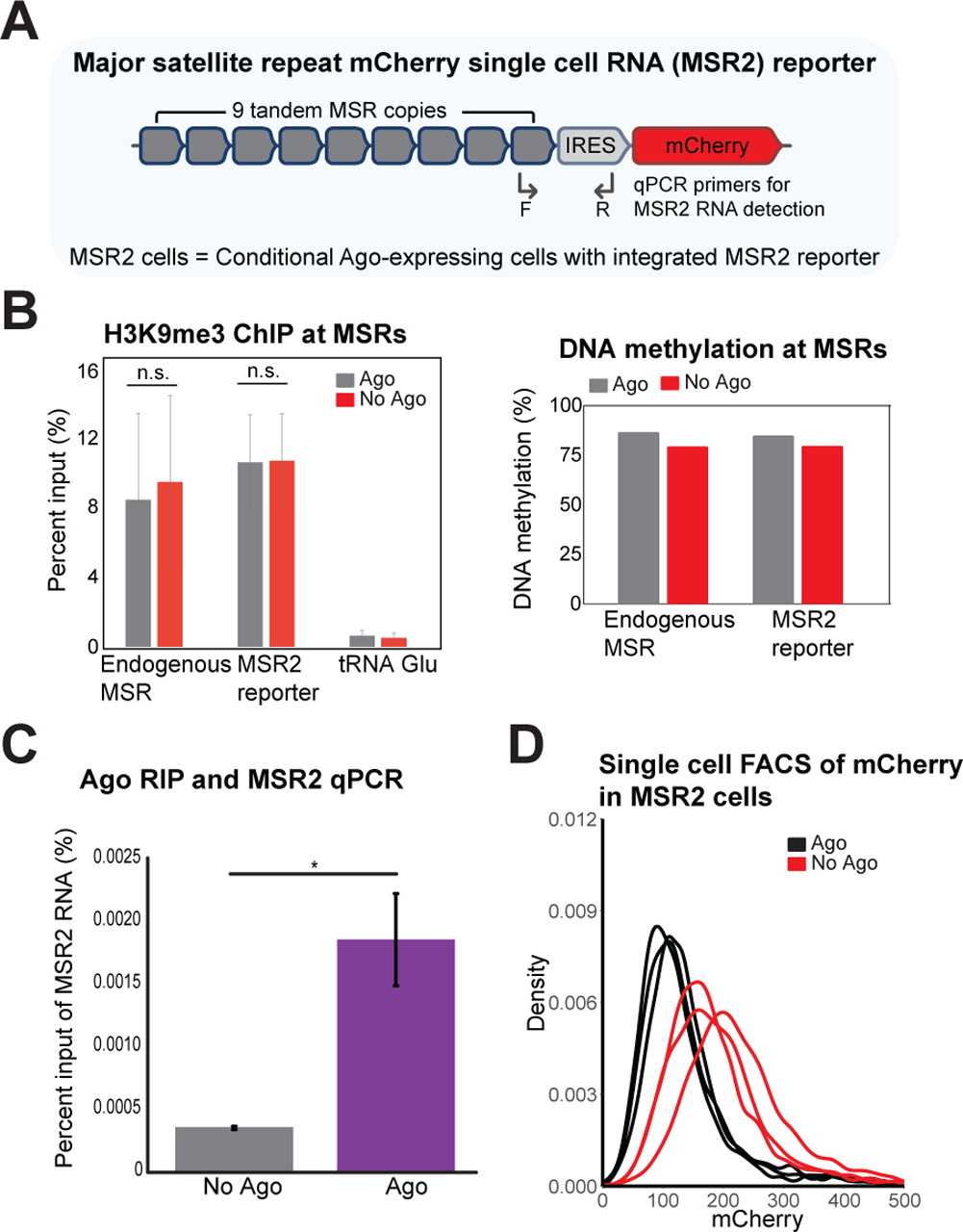
MSRs impart Ago-mediated regulation on an exogenous reporter. (A) Diagram of the MSR2 reporter containing and IRES-mCherry-pA cassette downstream of nine MSRs. Forward (F) and Reverse (R) primers for qPCR indicated. (B) Left: ChIP-qPCR for H3K9me3 at endogenous MSRs, the exogenous MSR2 reporter and a tRNA region as a negative control. ChIPs were performed in cells treated for Ago2 (Dox) or No Ago (No Dox) expression. Error bars show the standard error of biological triplicates. (n.s.: nonsignificant; by two-sided t-test of biological triplicates) Right: Barplot of the average DNA methylation determined by bisulfite DNA methylation assay. (C) Ago2 RIP-qPCR for the MSR2 reporter in cells treated for Ago2 (Dox) or No Ago (No Dox) expression. Error bars show the standard error of biological triplicates. (*: p value < 0.05; p value determined by two-sided t-test of biological triplicates) (D) FACS quantification of mCherry levels in MSR2 reporter cells treated for Ago2 (Dox) expression in black or No Ago (No Dox) expression in red. Biological triplicates shown.

We determined that the MSR2 reporter was targeted for heterochromatin formation since its histone H3 lysine 9 trimethylation (H3K9me3) and DNA methylation levels were similar to that of endogenous pericentromeres (Fig. 3B). Neither of these heterochromatin-associated modifications were significantly altered with Ago depletion at either endogenous or exogenous MSRs. Ago RIP followed by qPCR of the MSR2 reporter showed Ago-bound transcripts and confirmed RNAi targeting (Fig. 3C). To quantify MSR2 reporter expression, we used single-cell FACS measurements and found that mCherry levels increased upon Ago depletion, consistent with targeted repression by RNAi (Fig. 3D). These results indicate that as few as 9 tandem MSRs, which undergo heterochromatin formation similar to pericentromeres, are sufficient for Ago recruitment and repression.

### RNAi pericentromeric lncRNA repression occurs after S phase transcription

Previous studies found that pericentromeric RNA transcription occurs during DNA synthesis (Chen et al. 2008). The S phase-specific signal likely offers access for transcriptional machinery otherwise hindered by constitutive heterochromatin. We assessed MSR2 reporter activity in single cells and at different stages of the cell cycle to understand precisely when RNAi regulation occurs.

We established a four-color FACS assay detecting DNA levels (DAPI), S phase cells (BrdU incorporation), mitotic cells (histone H3 serine 10 phosphorylation, H3Ser10p) and MSR2 reporter levels (mCherry) (Fig. 4A). With this approach, we quantified MSR2 activity at specific cell cycle stages with and without Ago expression. We observed a pattern of increased MSR2 activity in late S phase and G2/M phase cells, compared to G1 phase and early S phase cells that occurred with or without Ago activity (Fig. 4B). These cell cycle differences are consistent with chromatin accessibility for transcriptional machinery near times of pericentromere DNA replication. The increase in MSR expression during S phase progression was validated for endogenous bidirectional pericentromeric lncRNAs in cell cycle synchronized populations (Supplemental Fig. S4A). The MSR2 reporter quantification confirmed increased MSR expression in all cell cycle stages in Ago-deficient cells (Fig. 4C; Supplemental Fig. S4B). Additionally, wider distributions of MSR2 activity in late S and G2/M phases in Ago-deficient cells indicate increased suppression after S phase transcription.

**Figure 4.**
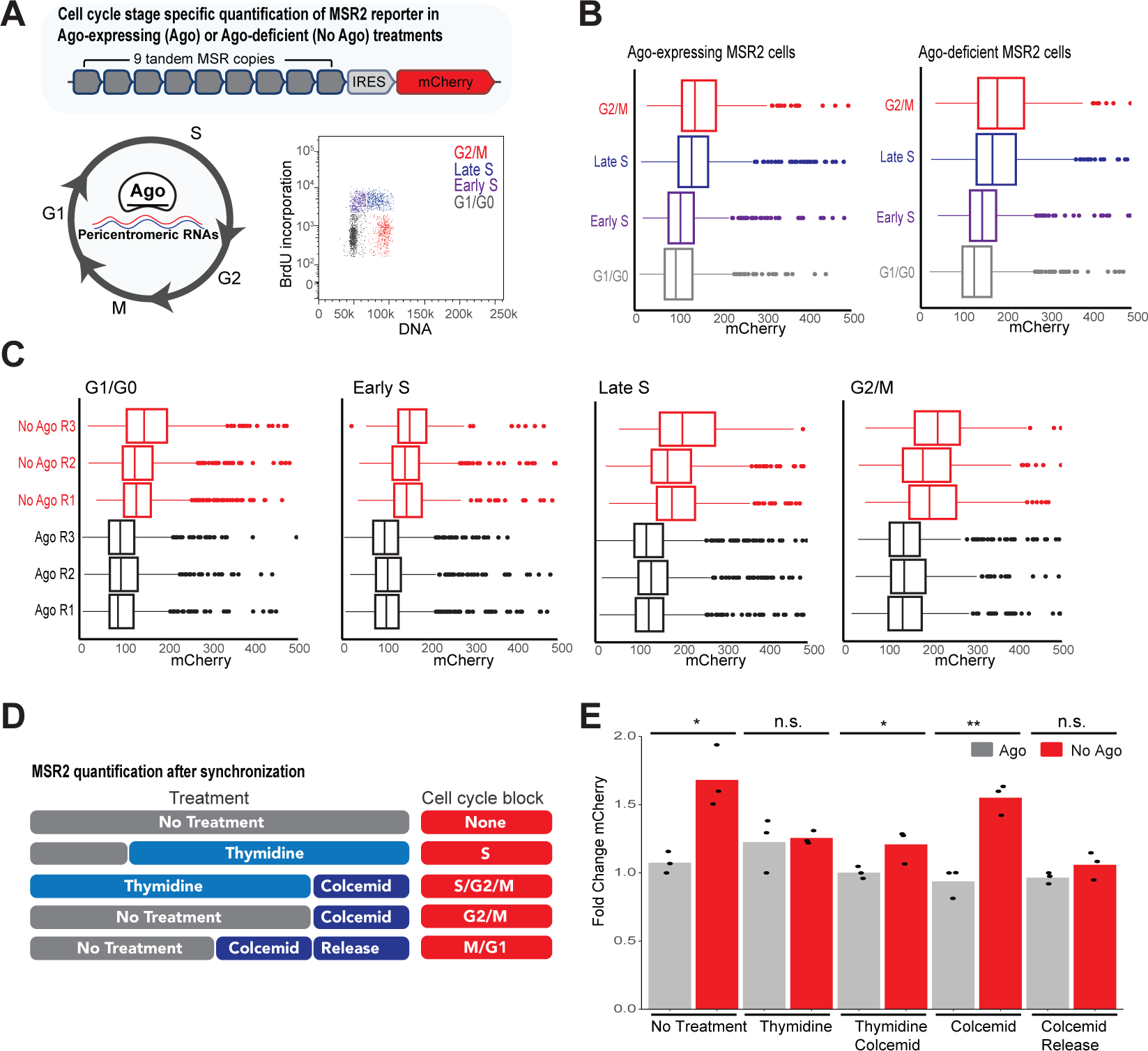
RNAi pericentromeric lncRNA repression occurs after S phase transcription. (A) Diagram of cell cycle stage characterization of MSR2 reporter activity to determine timing of RNAi regulation. Classification of cell cycle stages G0/G1, Early S, Late S and G2/M stages indicated. mCherry expression was quantified in gated populations. (B) FACS quantification of mCherry levels in MSR2 cells treated for Ago2 (Dox) or No Ago (No Dox) expression separated by cell cycle stages. (C) FACS quantification of mCherry levels at indicated cell cycle stage in MSR2 cells treated for Ago2 (Dox) or No Ago (No Dox) expression. Shown in biological triplicate. (D) Diagram of cell cycle treatments used to assay for cell cycle stage regulation. (E) Barplot for the average MSR2 activity in cells treated for Ago2 (Dox) expression in gray or No Ago (No Dox) expression in red. Cell cycle treatment indicated below. Expression normalized to Ago2-expressing biological replicate 1. (*: p value < 0.05; **: p value < 0.01; n.s.: nonsignificant; p value determined by two-sided t-test of biological triplicates)

We used cell cycle synchronization to confirm the timing of Ago regulation (Fig. 4D,E; Supplemental Fig. S4C-I). We found that early S phase synchronization decreased the difference of MSR2 expression between Ago-expressing and Ago-deficient cells, indicating an RNAi-independent turnover mechanism that functions near early S phase. Release from S phase into metaphase block resulted in increased MSR2 expression in Ago-deficient cells compared to Ago-expressing cells. This result further supports Ago-mediated repression after S phase transcription. The differences in MSR2 reporter activity in Ago-deficient cells were detected with metaphase block alone and decreased upon metaphase block release. These results are consistent with a role for Ago in suppressing pericentromeric lncRNAs after late S phase.

### Chromatin-associated pericentromeric lncRNAs are regulated by nuclear Ago

The MSR2 reporter activity indicates that Ago represses pericentromeric lncRNAs in the nucleus of interphase cells after S phase. To further support a nuclear Ago function, we aimed to colocalize Ago with pericentromere regions in interphase cells. The majority of Ago protein is cytoplasmic in proliferating cells and a small fraction is detected in the nucleus. We reasoned that if Ago was regulating pericentromeres after S phase, then we could use live-cell imaging to track Ago proximity to chromocenters over time. Chromocenters are perinuclear regions of compact pericentromeric heterochromatin that are clearly visible by dense DNA or histone staining in interphase cells. To track these regions in live cells, we generated mESCs that conditionally express 3x-mCherry-Ago2 and constitutively express H2B-GFP for colocalization with dense chromatin regions (Fig. 5A; Supplemental Movie 1). We generated high-resolution 3D time courses of mitosis in these cells (Supplemental Fig. S5A). The AI-determined signal over background showed relatively low levels of nuclear Ago compared to the cytoplasm as expected. For the nuclear Ago signal detected, we examined the overlap with dense chromatin regions (Fig. 5B). We found nuclear Ago signal was not only proximal to chromocenters at multiple z-planes within the nucleoplasm but tracked with chromocenter movement at multiple time points (Supplemental Fig. S5B). As expected, nuclear membrane breakdown during mitosis left chromosomes fully accessible to Ago interactions. We concluded that Ago has access to pericentromeric regions in interphase cells and next searched for additional support of nuclear Ago regulation.

**Figure 5.**
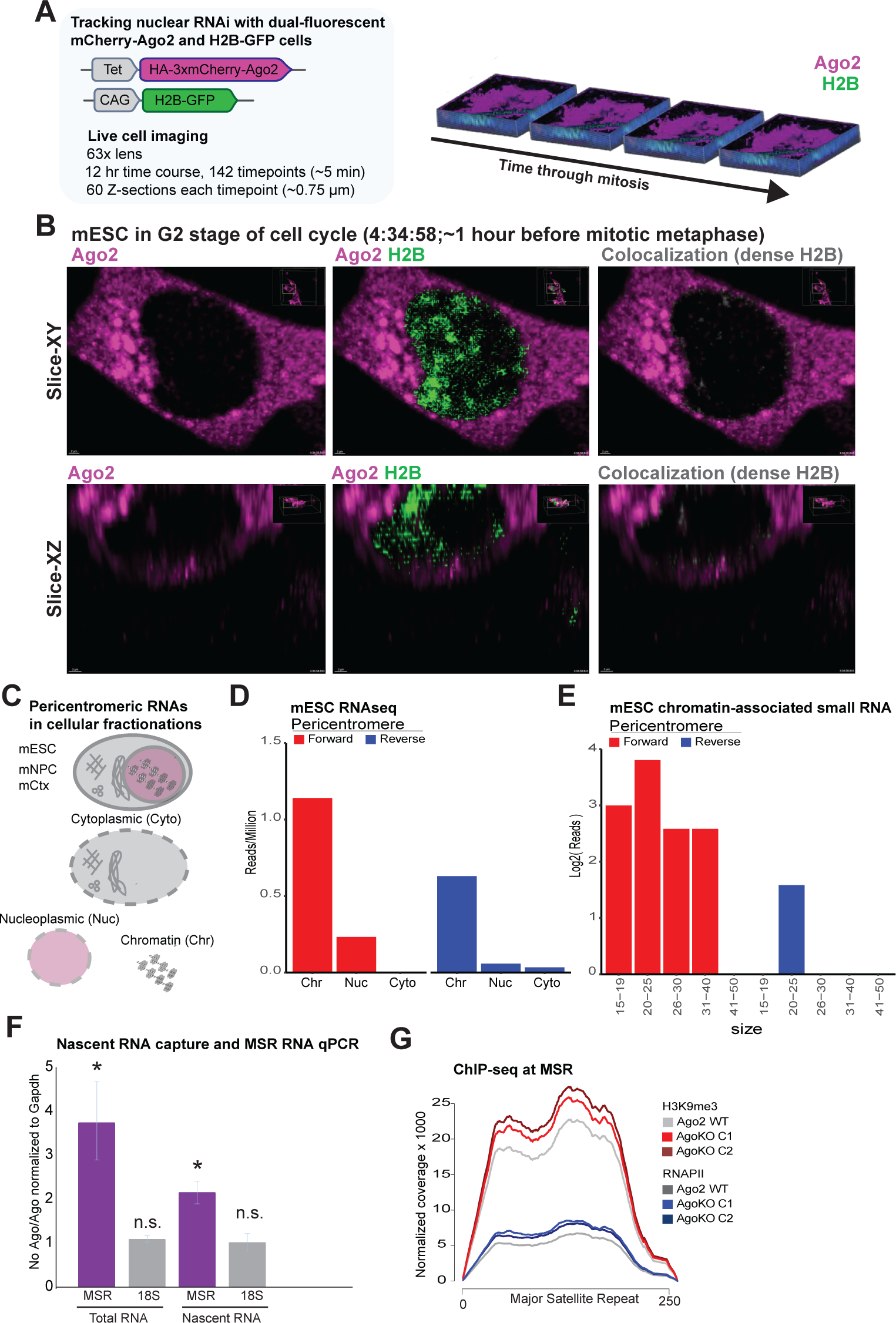
Chromatin-associated pericentromeric lncRNAs are regulated by nuclear Ago. (A) Diagram summarizing live cell imaging time course for HA-3xmCherry-Ago2 and H2B-GFP dual-fluorescent cells. (B) Colocalization of Ago2 with dense H2B-GFP heterochromatin in G2 phase of cell cycle. (C) Diagram of cellular fractionation in mouse embryonic stem cells (mESC), neuronal progenitor cells (mNPC) and primary cortical neurons (mCtx). Data from *Yeom et al*. (D) MSR counts from mESC cellular fractionation followed by total RNAseq. (E) MSR counts from mESC chromatin-associated small RNAseq. Reads were categorized by transcript orientation and RNA length. (F) Quantification of MSR and 18S nascent transcription in cells treated for No Ago (No Dox) expression compared to those treated for Ago2 (Dox) expression and normalized to Gapdh. (*: p value < 0.05; n.s.: nonsignificant; p value determined by two-sided t-test of biological triplicates) (G) ChIP-seq quantification of RNAPII and H3K9me3 levels at MSRs in Ago2 WT cells and in two clones (C1 and C2) of Ago KO cells.

Previous studies have indicated that pericentromeric RNAs are retained at pericentromeres (Lu and Gilbert 2007) (Probst et al. 2010). To determine the location of pericentromeric lncRNA regulation, we quantified pericentromeric RNAs in long and short RNAseq libraries from chromatin, nucleoplasm, and cytoplasm fractions (Yeom et al. 2021) (Fig. 5C). The long total RNAseq libraries revealed that bidirectional pericentromeric lncRNAs are enriched at chromatin in mESCs, neuronal progenitor cells (mNPCs) and primary cortical neurons (mCtxs) (Fig. 5D, Supplemental Fig. S5C). We also found that 20-25mers were enriched in the chromatin fractions in all three cell types suggesting local repression by Ago (Fig. 5E; Supplemental Fig. S5D).

To determine if Ago affects pericentromere transcription, we used 5-ethynyl uridine (EU) pulse labeling for nascent RNA capture, and measured RNA changes by RT-qPCR in conditional Ago-expressing cells (Fig. 5F). We found increased pericentromeric lncRNA transcription accounted for nearly half of the upregulation upon Ago loss, supporting both transcriptional and post-transcriptional repression. This result was supported by chromatin immunoprecipitation followed by DNA sequencing (ChIP-seq) data, in which we found RNA polymerase II (RNAPII) signal increased at MSRs with Ago loss. This increase in RNAPII was accompanied by similar increases in H3K9me3 in the ChIP-seq data (Fig. 5G). Together, these results support RNAi regulation at pericentromeres in interphase cells.

### RNAi loss results in mitotic defects that introduce genomic instability

Since dysregulation of pericentromeric lncRNA is linked to mitotic defects and genomic instability, we tracked cellular division with live-cell imaging of H2B-GFP in conditional Ago-expressing cells (Fig. 6A, Supplemental Movie 2,3). We observed severe mitotic defects in Ago-deficient cells that were characterized by progressive dispersion of H2B-GFP from metaphase-aligned chromosomes, as well as lagging H2B signal during anaphase leading to micronuclei formation (Fig. 6B). These abnormal mitotic characteristics were observed in nearly a quarter of all division events in Ago-deficient cells (Fig. 6C). We also determined general delays in mitosis in cells with otherwise normal mitotic H2B distributions (Fig. 6D), and a larger proportion of Ago-deficient cells spent nearly twice as long in mitosis compared to Ago-expressing cells (Fig. 6E; Supplemental Fig. S6A).

**Figure 6.**
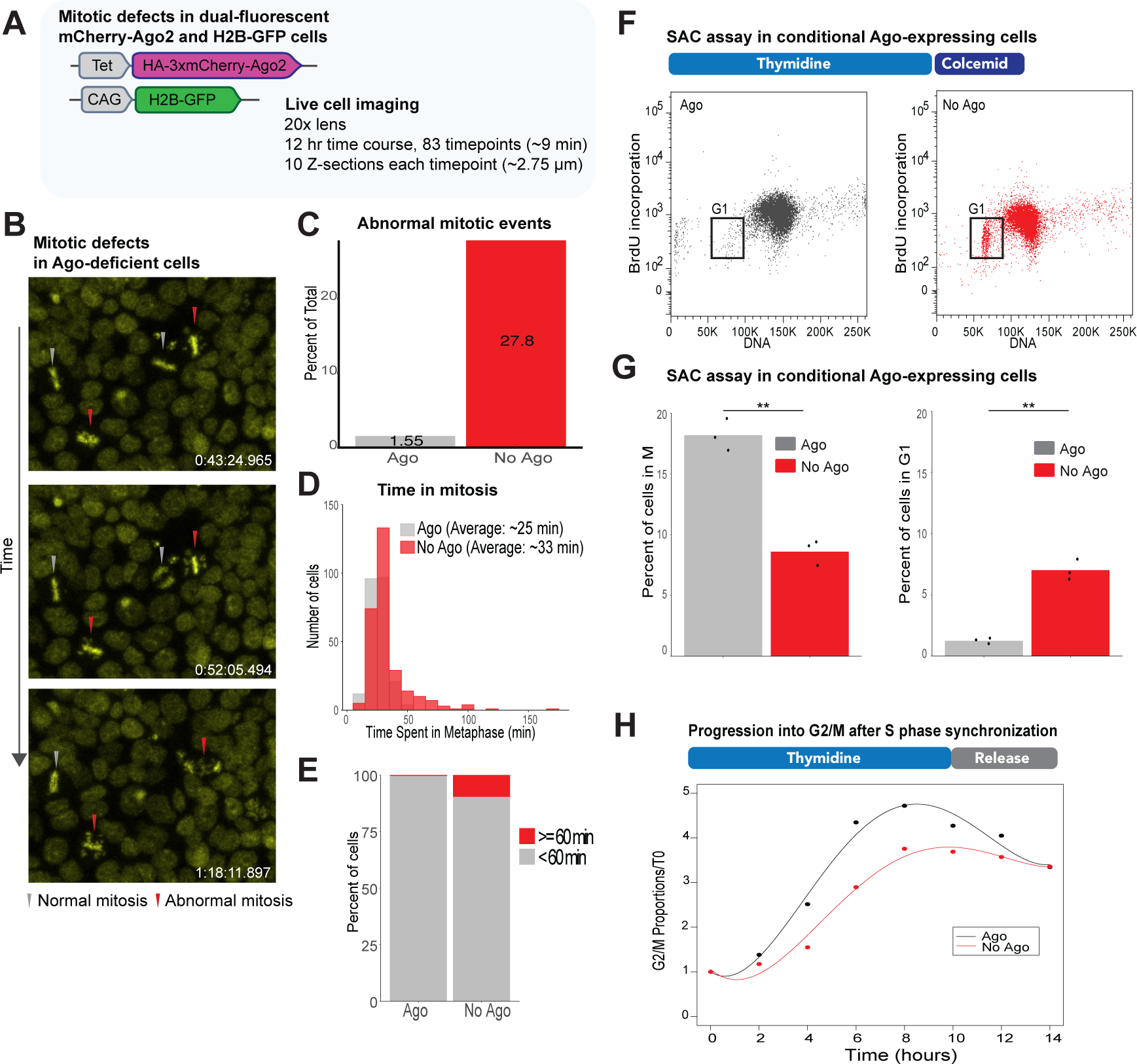
RNAi loss results in mitotic defects that introduce genomic instability. (A) Diagram of imaging conditions for HA-3xmcherry-Ago2 and H2B-GFP dual-fluorescent cells to track genomic instability. (B) Images from the time course of Ago-deficient cells with arrow heads indicating normal (gray) and abnormal (red) progression through mitosis. H2B-GFP shown in yellow. (C) Quantification of abnormal mitotic events from time course of cells treated for Ago2 (Dox) or No Ago (No Dox) expression. Percentage from: Ago2 = 194 mitotic events; No Ago = 205 mitotic events. (D) Quantification of time spent in metaphase determined from the time course of cells treated for Ago2 (Dox) or No Ago (No Dox) expression. Averages from: Ago = 231 mitotic events; No Ago = 282 mitotic events. (p value < 0.0001; p value determined by two-sided t-test) (E) Barplot showing the proportion of cells that spend > 60 min in metaphase in cells treated for Ago2 (Dox) or No Ago (No Dox) expression. (F) Scatterplot of cell cycle profiles in SAC assay in cells treated for Ago2 (Dox) or No Ago (No Dox) expression. Cells were stained for BrdU incorporation and DNA, with G1 population indicated. (G) Barplot for the average percent of cells in M phase (left) and G1/G0 (right) in the SAC assay. Percent normalized to Ago2 (Dox) expressing biological replicate 1. (**: p value < 0.01; p value determined by two-sided t-test of biological triplicates) (H) Plot of the proportion of G2/M cells after release from S phase block and measurement every 2 hours for 14 hours. Cell cycle progression for cells treated for Ago2 (Dox) or No Ago (No Dox) expression indicated. Lines are fitted using fourth degree polynomial curves, with residuals > 0.93 for all trials.

Since these mitotic defects were related to results previously linking pericentromeric lncRNA overexpression to altered SAC activity, we next assayed this important mitotic checkpoint. We observed a striking defect in the SAC, where Ago-deficient cells bypassed the requirement of chromosome spindle attachment and progressed through mitosis at higher levels than Ago-expressing cells (Fig. 6F,G). To confirm this SAC bypass, we used live-cell imaging of H2B-GFP to validate the erroneous anaphase phenotype (Supplemental Fig. S6B,C). Finally, we noted an increase in H3Ser10p levels in Ago-deficient cells compared to Ago-expressing cells, suggesting altered mitotic signaling by H3Ser10-modifying Aurora kinases (Supplemental Fig. S6D).

We next tracked cell cycle progression in synchronized cells to identify additional defects related to pericentromeric lncRNA dysregulation in interphase cells. We selected thymidine treatments to minimize interference from characterized G1→S defects related to miRNA loss in mESCs (Supplemental Fig. S6E). Interestingly, the proportion of cells that progressed into G2 phase was significantly higher in Ago-expressing cells than in Ago-deficient cells after 6-hour release from S phase (Supplemental Fig. S6F). Longer time courses through S phase revealed that early differences in the progression into G2/M phase leveled out at later time points, and confirmed a delayed progression through G2/M phase in Ago-deficient cells (Fig. 6H; Supplemental Fig. S6G,H).

Complementation of Ago null cells with Ago1 indicated that this Ago activity is independent of direct Ago2 cleavage and likely acts redundantly (Supplemental Fig. S7A-C). Similarly, we observed defects in cell cycle progression, mitotic SAC activity and genome stability in RNAi-mutant adult mMSCs (Supplemental Fig. S7D-F). In the adult mMSCs, we observed frequent defects in cytokinesis likely permitted by the absence of a functional p53 response (data not shown). Together, the results indicate that RNAi mutants have abnormal cell cycle progression and chromosome segregation contributing to genomic instability, and these defects are directly linked to forced overexpression of pericentromeric lncRNAs.

### Dysregulation of pericentromeric lncRNA drives RNAi mutant cell cycle defects

To directly connect mitotic defects in RNAi-deficient mESCs to pericentromeric lncRNA dysregulation, we investigated the consequences of pericentromeric lncRNA overexpression alone. First, we used dCas9-mediated transcriptional activation with dCas9-VP64 in MSR2 cells to confirm its ability to modulate MSR transcription (Fig. 7A,B). We found that targeted transcriptional induction was able to specifically increase MSR2 activity and was sufficient to increase H3Ser10p levels in mitotic cells compared to nontargeted controls.

**Figure 7.**
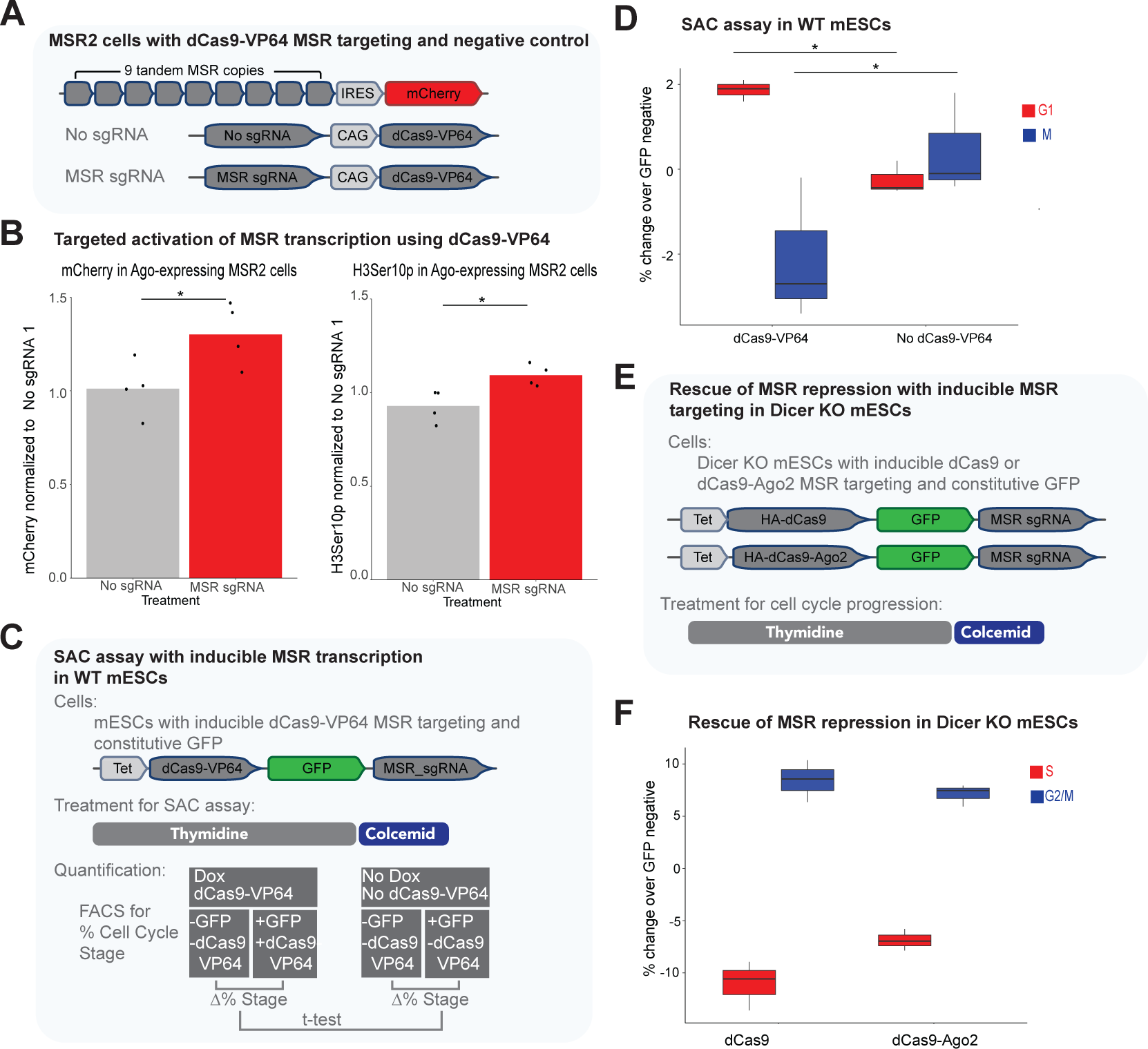
Dysregulation of pericentromeric lncRNA drives RNAi mutant cell cycle defects. (A) Diagram summarizing targeted MSR induction experiment. The constitutive expression of dCas9-VP64 with or without targeting to MSRs in MSR2 reporter cells. (B) Expression of dCas9-VP64 in MSR2 reporter cells treated for Ago2 (Dox) or No Ago (No Dox) expression and colcemid for enrichment of mitotic cells. Right: Barplot of the average mCherry expression. Left: Barplot of average H3Ser10p levels. Levels are normalized to Ago2 expressing replicate 1 without sgRNA treatment. (*: p value < 0.05; p value determined by two-sided t-test with four biological replicates) (C) Diagram for SAC activity assay in WT mESCs with inducible MSR induction. Comparison of dCas9-VP64 targeting in mixed population of integrated (+GFP) and nonintegrated (-GFP) cells. (D) SAC activity in WT mESCs with MSR transcription induction. Quantification of the percent change in G1/G0 (red) and M (blue) phase cells following S phase synchronization and release into metaphase block with colcemid. (*: p value < 0.05; p value determined by two-sided t-test of biological triplicates) (E) Diagram of MSR repression by transcriptional interference or local recruitment of Ago. (F) Quantification of percent change in S (red) and G2/M (blue) cell cycle populations following S phase synchronization and release into colcemid in Dicer KO mESCs. Dox treatments for the expression of either dCas9 or dCas9-Ago2 in mixed population of integrated (+GFP) and nonintegrated (-GFP) cells. (differences between −GFP and +GFP population at all four stages/treatments have p values < 0.01; p value determined by two-sided t-test of biological triplicates)

After confirming MSR transcriptional modulation, we quantified cell cycle progression defects. We found that the inducible expression of dCas9-VP64 within a mixed population of stably integrated (+GFP) and nonintegrated (-GFP) cells allowed the most direct comparison (Fig. 7C). With this approach, we determined that the induction of MSR transcription in WT mESCs was sufficient to cause the SAC defect (Fig. 7D), directly linking MSR overexpression to mitotic defects in mESCs. Next, we rescued the cell cycle progression defects in Dicer KO mESCs by targeting dCas9 or dCas9-Ago2 fusions to MSRs (Fig. 7E,F). We found that either dCas9 or dCas9-Ago2 targeting was sufficient to rescue progression through S phase and increase the accumulation of cells in G2/M phase. These results indicate that dCas9-mediated transcriptional interference at MSRs alone can rescue cell cycle progression defects of Dicer KOs, and likely accounts for the dCas9-Ago2 rescue activity.

## Discussion

In this study, we show that RNAi functions in genome stability by suppressing bidirectionally transcribed pericentromeric lncRNAs. We determine molecular characteristics of pericentromeric lncRNAs and their uniform upregulation upon loss of RNAi. We present evidence supporting repression by nuclear Ago in interphase cells after pericentromeric RNA transcription in S phase. We link the increased pericentromeric lncRNA levels in RNAi mutant mESCs to genomic instability. The observations of this RNAi activity in two somatic cell types supports a general role in cell division.

Several factors contribute to conflicting studies of RNAi activity at mammalian pericentromeres (Murchison et al. 2005; Kanellopoulou et al. 2005; Fukagawa et al. 2004). Recent characterization in Dicer KO mESCs highlighted genotype differences in clonal knockout lines selected for during adaptation to RNAi loss (Gutbrod et al. 2022). Here, we focused on conditional expression of Ago showing reversible regulation in a single cell line. We characterize pericentromeric lncRNAs in the presence or absence of Dicer and with rescue by multiple Ago proteins. Focusing this study on the Ago proteins, the effectors of the RNAi pathway, allowed the identification of a direct function for RNAi at pericentromeres.

In fission yeast, autoregulation by degradation intermediates from primary transcripts is well-characterized (Yamanaka et al. 2013) (Halic and Moazed 2010). Similar to these autoregulatory guides in fission yeast, mammalian pericentromeric small RNA guides are detected at low copy number. Classification through both Ago- and Dicer-dependence aids in separating lowly expressed guides from other degradation intermediates. The retention of these small RNAs and target lncRNAs near chromatin likely enhances the Ago-mediated regulation. Phase transitions at pericentromeres (Larson et al. 2017; Henninger et al. 2021) may also contribute to higher local concentrations of functional and regulatory RNAs in this region.

The mapping of Ago-dependent pericentromeric small RNAs on the MSR indicates a positional correlation between the forward strand and hairpin regions from the reverse strand. These hairpins may act to slow degradation allowing for the formation of short dsRNA Dicer substrates. A reliance on degradation intermediates as Dicer substrates, as opposed to long dsRNA precursors, may avoid dsRNA-activated immune responses in mammalian cells. Expanded detection and classification of autoregulatory small RNAs from other degradation fragments will be impactful in addressing Ago-mediated suppression at chromatin genome-wide.

Transcriptional activity is likely limited to a relatively small subset of MSRs based on the sizes we determined for the pericentromeric lncRNAs. Considering the distinct pericentromeric lncRNA lengths revealed in the RNase H assay, we propose that they are cleaved and protected from degradation for a period of time to allow for a local function. These bidirectional lncRNAs may be kept from annealing to each other by the binding of protein complexes near pericentromeres. These pericentromeric lncRNAs are transcribed by RNAPII, but their 5’ termini indicate processing events that are distinct from mRNA and may contribute to chromatin retention and local activity. Determining RNA processing and degradation factors that act on these lncRNAs will be important in characterizing their full function during cell division.

We propose that mammalian RNAi activity at pericentromeres serves to suppress functional pericentromeric lncRNAs. The characterization of all mammalian RNAi mutants to date indicates that the direct link between RNAi and heterochromatin factor recruitment differs substantially from other model systems. An adaptation of mammalian RNAi to the regulation of functional pericentromeric lncRNAs may allow expanded nuclear Ago functions. Pericentromeric lncRNAs could contribute directly to cell cycle progression by impacting the efficiency of heterochromatin formation (Burton et al. 2020) (Novo et al. 2022), or the RNA-induced activity of cell cycle regulated kinases (Jambhekar et al. 2014; Blower 2016).

Uncovering the mechanistic links between these noncoding RNAs and chromosomal behavior will advance our understanding of genome stability and noncoding RNA defects that contribute to cancer.

## Materials and methods

### Cell culture

mESCs were grown on gelatinized tissue culture plates in Dulbecco’s Modified Essential Media supplemented with HEPES (Thermo Fisher), 15% fetal bovine serum (Hyclone/Thermo Fisher), 1000 U/mL leukemia inhibitory factor (Millipore), 0.1 mM non-essential amino acids (Gibco), 0.1 mM L-glutamine (Gibco), 0.1 mM penicillin/streptomycin (Gibco) and 0.11 mM β-mercaptoethanol (Gibco). mMSCs were grown in α-MEM (Gibco) supplemented with 10% FBS and 0.1 mM penicillin/streptomycin (Gibco). All cells were cultured with 5% CO2 and ambient oxygen at 37 ℃. Conditional Ago-expressing mESCs were maintained in 0.1 µg/ml doxycycline (Dox) (Sigma). For induction to wildtype Ago levels, cells were cultured in 2.5 µg/ml Dox for 48 hours. For depletion, cells were grown without Dox for 96 hours.

### Generation of transgenic cell lines

Dicer knockouts were generated using two Cas9 plasmids (px330) targeting regions flanking Dicer RNase III domains. These two plasmids and a plasmid conferring puromycin resistance were transfected using Lipofectamine^TM^ 3000 (Invitrogen). Cells were selected with 1 μg/ml puromycin (Gibco) after 24 hours. After 48-hour selection, puromycin-resistant cells were diluted into 96-well plates for growth of colonies from single cells. The colonies were expanded and genotyped by PCR.

For stable expression of HA-3xmCherry-Ago2, H2B-GFP, MSR2 reporter, and Dox-inducible dCas9 targeting, constructs were integrated using transient transfection with mPBase (piggybac transposase expression plasmid, a gift from A.W. Cheng).

### Northern blotting

For low-resolution northerns, 5 µg of total RNA was resolved on a 1.2% formaldehyde agarose gel and transferred by capillary to a Hybond-NX (Amersham) membrane. For medium-resolution northerns, 5 µg of total RNA was resolved on an 8% polyacrylamide urea gel and transferred to a HyBond-NX membrane (Amersham) using a TransBlot SD Semi-dry Transfer System (Bio Rad).

RNA was crosslinked twice using optimal crosslink setting on a Spectrolinker XL-1000. Crosslinked membranes were blocked in ULTRAHyb (Invitrogen) for random hexamer probes or oligonucleotide pre-hyb (6X SSC, 1X Denheardt’s solution, 0.5%SDS, 10 µg/ml yeast tRNA) for 30 minutes at 42 ℃ with rotation. Membranes were hybridized with radiolabeled probes overnight at 42 ℃ with rotation. Random hexamer probed membranes were rinsed twice (2X SSC and 0.1% SDS buffer) at 23 ℃, and then washed twice (0.2X SSC and 0.1% SDS buffer) for 30 minutes at 42 ℃ with rotation. Oligonucleotide-probed membranes were rinsed twice (2X SSC and 0.1% SDS buffer) at 23 ℃, and then washed twice (2X SSC and 0.1% SDS buffer) for 30 minutes at 42 ℃ with rotation.

MSR random hexamer probes were generated using the SalI/NotI fragment from the pγsat plasmid. The pγsat was a gift from Niall Dillon (Addgene plasmid # 39238; http://n2t.net/addgene:39238; RRID:Addgene_39238). The Gapdh random hexamer probe was generated using primers described in Table S1 for qPCR. Oligonucleotides used for northern probes are listed in Table S1. The blots were exposed to phosphor screens (GE HealthCare) for visualization. Images were adjusted for contrast using ImageJ (NIH) software and cropped for presentation in Illustrator (Adobe).

### RNA isolation and quantitative RT-PCR

Cells were washed once in 1X HEPES-buffered saline (HBS) and TRIzol (Life Technologies) added directly to culture plates. Total RNA was extracted from TRIzol using manufacturer instructions. For removal of contaminating MSR DNA, extensive DNase treatment with TURBO DNaseI (Life Technologies) required that 10 µg of total RNA was incubated twice with 2 U TURBO DNaseI for 30 minutes at 37 ℃. RNA was purified again after DNase treatment using acid phenol chloroform (Ambion). 1 µg of RNA was reverse transcribed using Superscript IV (Life Technologies) with random hexamer primers. cDNA was amplified with PowerTrack SYBR Green Master Mix (Applied Biosystems) on a LightCycler 480 System (Roche). Primers used are listed in Table S1.

### RNase H, Xrn1, polyA pulldowns and RACE

10 µg of DNase-treated total RNA was digested with 5 U of RNase H (NEB) and 2 picomoles of a chimeric oligonucleotide targeting the MSRs for 20 minutes at 37 ℃. The Xrn1 (NEB) treatments were performed with 1 U of enzyme and incubation for 1 hr at 30 ℃. For polyA pulldown, 10 µg of DNase-treated total RNA was incubated with Dynabeads Oligo (dT)_25_ following manufacturer instructions. For rapid amplification of cDNA ends (RACE), 5 µg RNase H-treated RNA was poly-A tailed with 2 U of poly(A) polymerase (Takara) for 1 hour at 37 ℃. After acid-phenol purification, 800 ng RNA was used with the SMARTer RACE 5’/3’ kit (Takara) following manufacturer instructions. Oligonucleotides used for RNase H cleavage and RACE experiments are listed in Table S1.

### Cell cycle profiling and intracellular staining

For S phase synchronization, cells were grown in 300 µg/ml thymidine (Sigma). For metaphase block, cells were treated with KaryoMAX colcemid solution (Gibco) at 200 ng/ml. To label S phase cells, BrdU from the BD Pharmingen BrdU Flow Kit (BD Biosciences) was used at 1mM final concentration for 40 minutes. At the time of collection, cells were washed in 1X HBS and trypsinized. Cell pellets were resuspended in 1X HBS and fixed with 2% paraformaldehyde. All samples were analyzed on the LSRFortessa Cell Analyzer (BD Biosciences). Data were collected using FACS Diva (BD Biosciences) software and data was analyzed using FlowJo and custom R scripts.

### Nascent transcription quantification

To capture nascent RNA, the Click-iT Nascent RNA Capture Kit (Life Technologies) was used following manufacturer instructions. mESCs were pulsed with 0.5 mM 5-ethynyl uridine (EU) for 1 hour. 5 µg of purified total RNA was biotinylated with 0.5 mM biotin-azide and purified by ethanol precipitation. 1 µg of biotinylated RNA was used for pulldown. Reverse transcription with Superscript IV (Life Technologies) was performed on-bead.

### Chromatin immunoprecipitation

ChIP was performed as described previously(Marson et al. 2008). Approximately 30 million cells were crosslinked for 15 min at room temperature in 1% formaldehyde (Pierce). Excess formaldehyde was quenched for 5 minutes with 125 mM glycine. For immunoprecipitations, Protein G Dynabeads (Pierce) were blocked with 0.5% BSA (w/v) in phosphate-buffered saline (PBS) solution and conjugated with anti-H3K9me3 (Abcam, ab8898) antibody. Beads were washed twice in Sonication buffer (20mM Tris-HCl, pH 8.0, 150mM NaCl, 2mM EDTA, 0.1% SDS, 1% Triton X-100), twice in Sonication buffer + 500mM NaCl (20mM Tris-HCl, pH 8.0, 500mM NaCl, 2mM EDTA, 0.1% SDS, 1% Triton X-100), LiCl wash (10mM Tris-HCl, pH 8.0, 250mM LiCl, 1mM EDTA, 1% NP-40), and TE + 50mM NaCl (10mM Tris-HCl, pH 8.0, 1mM EDTA, 1% NP-40). DNA was eluted, crosslinks were reversed and DNA was purified by phenol chloroform extraction and ethanol precipitation.

### RNA immunoprecipitations

Cell pellets were lysed in CLIP lysis buffer (1% NP-40, 0.1% SDS, 130 mM NaCl, 50 mM Tris pH 7.4, 1 mM EDTA, 1mM DTT, 60 U SuperRNaseIn (Invitrogen), 1X protease inhibitor cocktail (Pierce)). Cell extracts were rotated at 4 ℃ for 30 minutes to ensure lysis. Cell lysates were sonicated with a Bioruptor (Diagenode) to remove all viscosity (high setting, 6 cycles of 30 seconds on, 30 seconds rest). The cell lysate was then centrifuged at 20,000xg for 15 minutes at 4 ℃ and the supernatant was used for IP. FLAG M2 antibody (Sigma) was conjugated to protein G Dynabeads (Pierce) for 2 hours in 0.1 M Na-Phosphate pH 8.0 at 4 ℃ with rotation. Extracts were incubated with antibody overnight at 4 ℃ with rotation. Beads were collected and washed twice with 1X PBS, twice with High Salt wash buffer (1% NP-40, 0.1% SDS, 1 M NaCl, 50 mM Tris pH 7.5, 1 mM EDTA), and twice with Low Salt wash buffer (1% NP-40, 0.1% SDS, 300 mM NaCl, 50 mM Tris pH 7.5, 1 mM EDTA) for 10 minutes each with rotation at 4 ℃. Washed beads were placed directly in TRIzol for isolation of RNA.

### Live cell imaging

Cells were seeded at 2e^5^/chamber on glass bottom chamber slides (Ibidi) that were pretreated with StemXVivo Culture Matrix (R&D Systems). Cells grown on chamber slides were imaged on a Zeiss LSM 880 Confocal with an environmental chamber for temperature and CO_2_ control. Images were processed with Imaris software (v10.1) for deconvolution, background processing, colocalization and filter color.

### Plasmids

For generation of Dicer KOs, oligonucleotides were annealed and cloned into pX330 plasmid using BbsI sites. pX330-U6-Chimeric_BB-CBh-hSpCas9 was a gift from Feng Zhang (Addgene plasmid # 42230; http://n2t.net/addgene:42230; RRID:Addgene_42230). Repeat sequences from pγsat were assembled using the MXS-chaining strategy(Sladitschek and Neveu 2015) to generate the MSR2 reporter. Piggybac recombination sites were amplified from piggybac expression vector, PBNeoTetO-Dest (a gift from A.W. Cheng).

For the constitutive dCas9-VP64 expression vector, dCas9-VP64 from pAC147-pCR8-dCas9VP64 was cloned into pX335-U6-Chimeric_BB-CBh-hSpCas9n(D10A) using EcoRI/AgeI sites. pAC147-pCR8-dCas9VP64 was a gift from Rudolf Jaenisch (Addgene plasmid # 48219; http://n2t.net/addgene:48219; RRID:Addgene_48219) (Cheng et al. 2013). pX335-U6-Chimeric_BB-CBh-hSpCas9n(D10A) was a gift from Feng Zhang (Addgene plasmid # 42335; http://n2t.net/addgene:42335; RRID:Addgene_42335) (Cong et al. 2013). dCas9-Ago2 fusions were made from mouse Ago2 cDNA. Oligonucleotides used to generate plasmids are listed in Table S1.

### Bisulfite Sequencing

Genomic DNA was used with the EpiMark Bisulfite Conversion Kit (NEB) using the manufacturer instructions.

### RNA sequencing and analysis

For total RNA sequencing, rRNA was removed using the RiboMinus Eukaryotic Kit (Invitrogen) and library preparation performed using the TruSeq Stranded mRNA Library Prep kit (Illumina). Small RNA (Zamudio et al. 2014) and iCLIP (Bosson et al. 2014) reads were processed as described previously. Tandem repeats for major satellite (V00846.1) (Hörz and Altenburger 1981), minor satellite (X14462.1) (Wong and Rattner 1988), rDNA (BK000964.1) (Grozdanov et al. 2003) and telomere repeats were appended to the mm10 mouse genome build for mapping. Reads were mapped directly to the MSR consensus sequence using the Bowtie (Langmead et al. 2009) short read alignment tool allowing no mismatches. Reads were quantified using a combination of BEDTools (Quinlan and Hall 2010) commands and R commands for visualization.

### Quantification and statistical analysis

All statistical tests are described in figure legends.

### Data sets

The data sets generated and analyzed in this study will be available in NCBI GEO. The raw images will be available in a Mendeley data set.

## Acknowledgements

The authors thank the UCLA BSCRC Flow Cytometry and Microscopy Core facilities. We thank Tim Kelly, Jesse Martinez, Emma Breen, John Garcia, Nima Hooshdaran and Mito Tariveranmoshabad for technical support and helpful discussions. R.S. was supported by the Ruth L. Kirschstein NRSA (GM007185). This work was supported by the NIH-NIGMS (J.R.Z., 5R01GM143536) and the Kure It Cancer Research Foundation award (J.R.Z.).

## Author contributions

J.R.Z. conceived the study. R.S., V.P., N.R.G., and C.N.D. performed and analyzed experiments with the guidance of J.R.Z. R.S. performed RNA characterization of pericentromeric lncRNA. C.N.D. and V.P. analyzed live cell imaging data. R.S. and C.N.D. performed ChIP- and RIP-qPCR experiments. C.N.D. and N.R.G. analyzed FACS data for the characterization of cell cycle progression. The manuscript was written by R.S and J.R.Z with input from all authors.

