## Supplemental Figures for "Mammalian RNAi represses pericentromeric lncRNAs to maintain genome stability"

Sandoval_Figure_S1

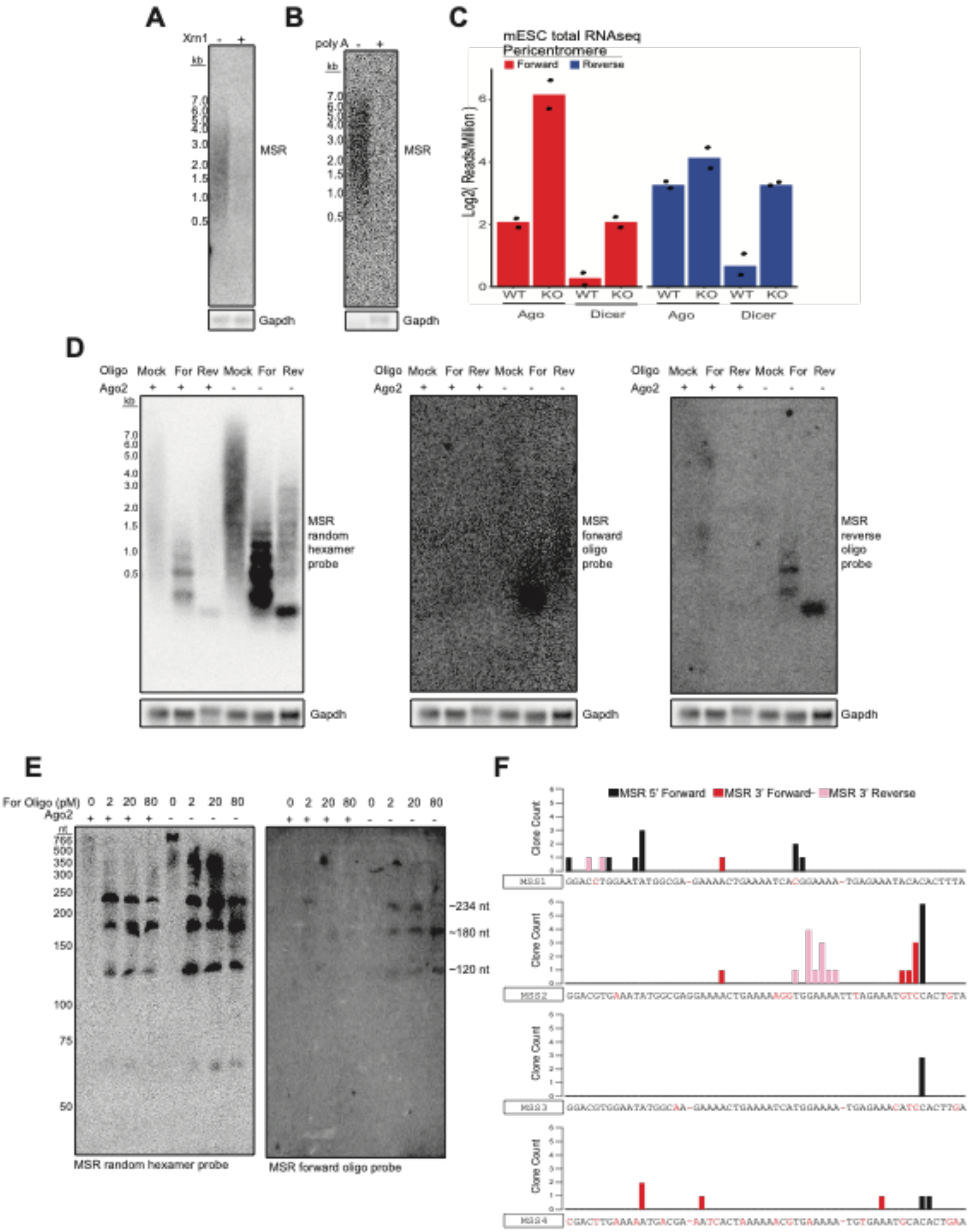

**Supplemental Figure S1. The RNAi pathway uniformly represses bidirectional pericentromeric lncRNAs**

(A) Northern blot of total RNA from cells treated for No Ago (No Dox) expression and after digestion with the 5'-phosphate-dependent 5'→3' exonuclease Xrn1 or mock treatment. Top: Random hexamer probe for MSRs; Bottom: Random hexamer probe for Gapdh.

(B) Northern blot of total RNA from cells treated for No Ago (No Dox) expression after selection for polyA sequences. +: positive selection or bound RNA, -: negative selection or unbound RNA. Top: Random hexamer probe for MSRs; Bottom: Random hexamer probe for Gapdh.

(C) Total RNAseq quantification of MSR reads in Ago1-4 WT and Ago1-4 KO mESCs, or Dicer WT and Dicer KO mESCs. Forward mapping reads plotted in red and reverse mapping reads plotted in blue.

(D) Reprobing of RNase H cleaved total RNA from cells treated for Ago (Dox) or No Ago (No Dox) expression. Left panel from Fig. 1E and is included for reference of signal from reprobing with oligonucleotides complementary to forward MSR transcripts (center) and reverse MSR transcripts (right).

(E) Medium resolution northern blot of total RNA from cells treated for Ago (Dox) or No Ago (No Dox) expression. Forward targeting chimeric oligonucleotide for RNase H cleavage was used at indicated concentrations. Left: MSR random hexamer probe; Right: MSR forward transcript oligonucleotide probe. The ~234 nt indicates the repeat unit fragment and smaller products that are likely the result of off-target cleavage at other MSSs.

(F) Barplot of Rapid Amplification of cDNA Ends sequencing results for forward and reverse transcripts in total RNA from cells treated for No Ago (No Dox) expression and cleaved with RNase H using the forward targeting chimeric oligonucleotide.

**Sandoval_Figure_S2**

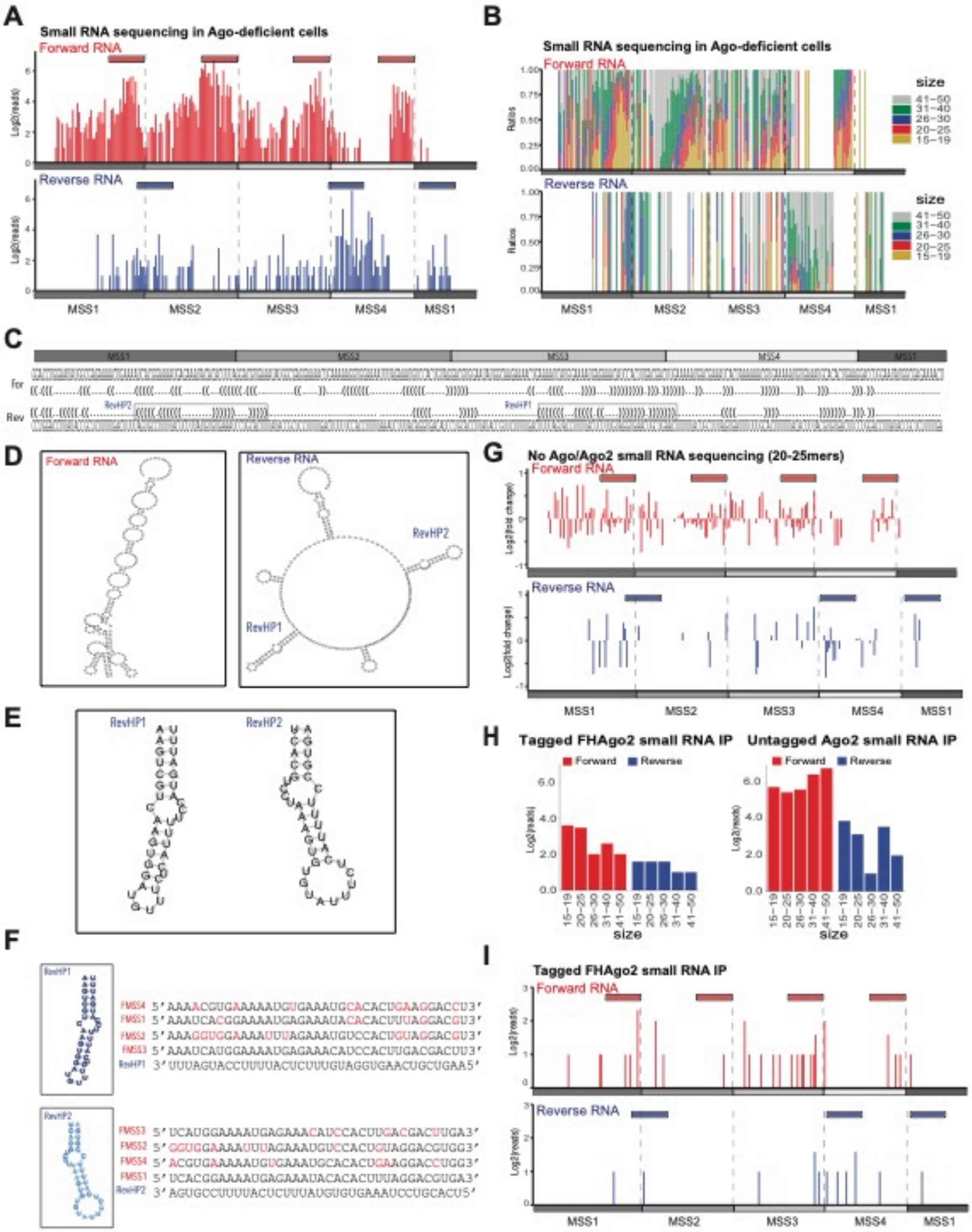

**Supplemental Figure S2. Ago- and Dicer-dependent autoregulatory small RNAs from pericentromeric lncRNAs**

(A) Small RNA profile in cells treated for No Ago (No Dox) expression. 5' RNA positions are indicated along the 234 nt MSR. Gray diagram below the plot indicates the subrepeats that include MSS1 of a second repeat on the left. Forward (red) and reverse (blue) reads are plotted separately with bars above the plot indicating the referenced clusters.

(B) Barplot of the proportion of indicated size ranges at each nucleotide position in cells treated for No Ago (No Dox) expression. Only the position of the 5' end of small RNA is plotted. Near the MSS 3' ends on the forward RNA staggered fragment lengths indicate 5'→3' degradation.

(C) Secondary structure prediction of the forward and reverse pericentromeric repeats. RNAfold was used with default settings. Positions of reverse hairpins overlapping forward degradation fragment clusters indicated.

(D) Diagram of predicted secondary structure for forward (left) and reverse (right) RNA.

(E) Diagram of two longest predicted hairpins in the reverse RNA.

(F) Reverse hairpins (RevHP) predicted in the pericentromeric RNA aligned to forward MSS1-4. Mismatched nucleotides between the forward MSS (FMSS) and RevHP indicated in red.

(G) The log2 fold change of the proportion of 20-25mers along the MSR in forward (red) and reverse (blue) orientation in cells treated for No Ago (No Dox) or Ago2 (Dox) expression. Plotted as described in panel A.

(H) Small RNA reads mapping to the pericentromeric forward (red) and reverse (blue) MSR grouped by size range. FLAG and HA epitope-tagged FHAgo2 small RNA IP on the left and negative control untagged Ago2 on the right.

(I) Small RNA profile from epitope-tagged FHAgo2 small RIP. Plotted as described in panel A.

Sandoval_Figure_S3

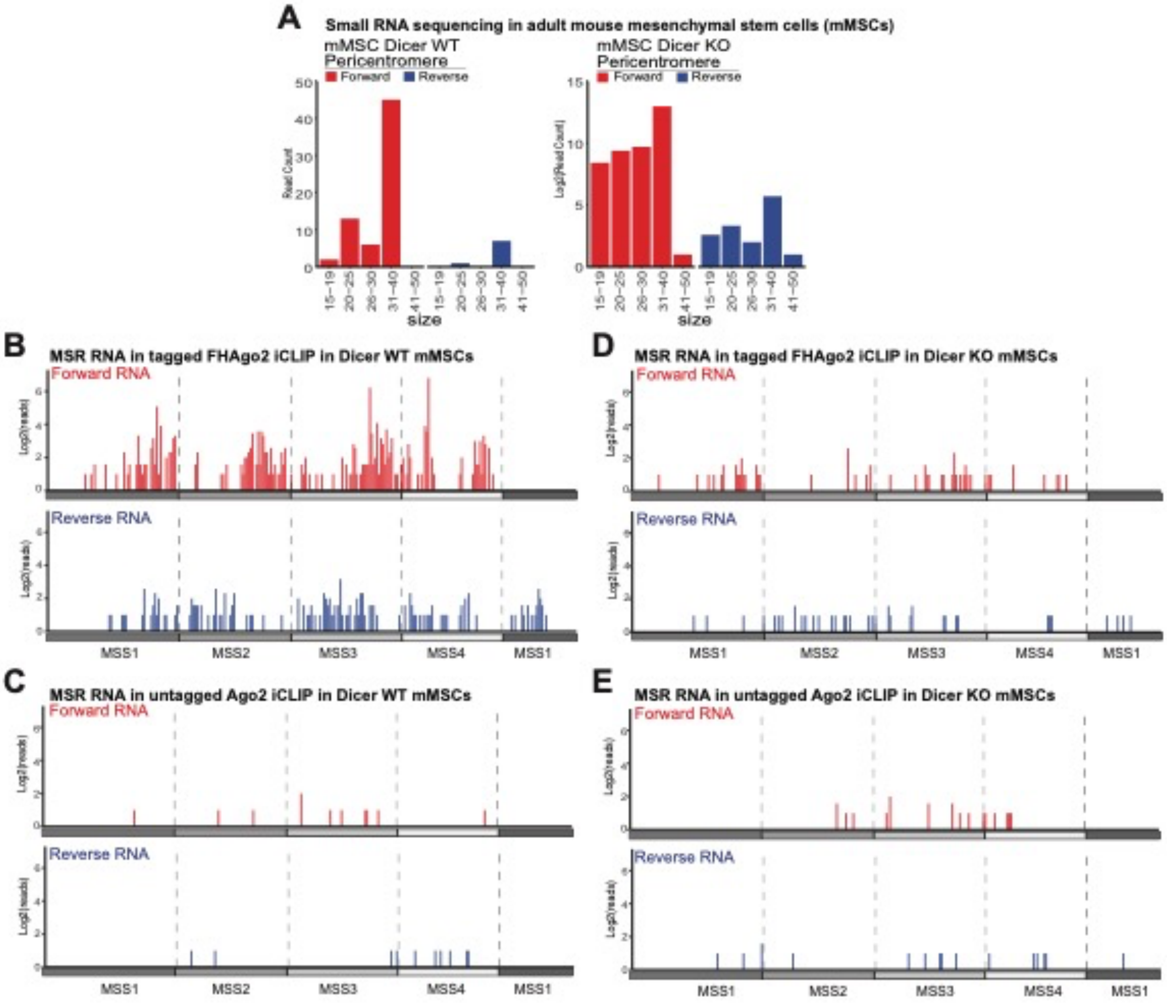

**Supplemental Figure S3. Dicer-dependent small RNA and Ago-pericentromeric lncRNA interactions in adult mesenchymal stem cells.**

(A) Quantification of small RNA reads in mMSCs mapped to forward (red) and reverse (blue) MSR categorized by size range in indicated Dicer genotypes.

(B) FLAG and HA epitope-tagged FHAgo2 iCLIP reads in mMSC mapping to forward (red) and reverse (blue) MSR in Dicer WT cells.

(C) Negative control untagged Ago2 iCLIP reads in mMSC mapping to forward (red) and reverse (blue) MSR in Dicer WT cells.

(D) FLAG and HA epitope-tagged FHAgo2 iCLIP reads in mMSC mapping to forward (red) and reverse (blue) MSR in Dicer KO cells.

(E) Negative control untagged Ago2 iCLIP reads in mMSC mapping to forward (red) and reverse (blue) MSR in Dicer KO cells.

**Sandoval_Figure_S4**

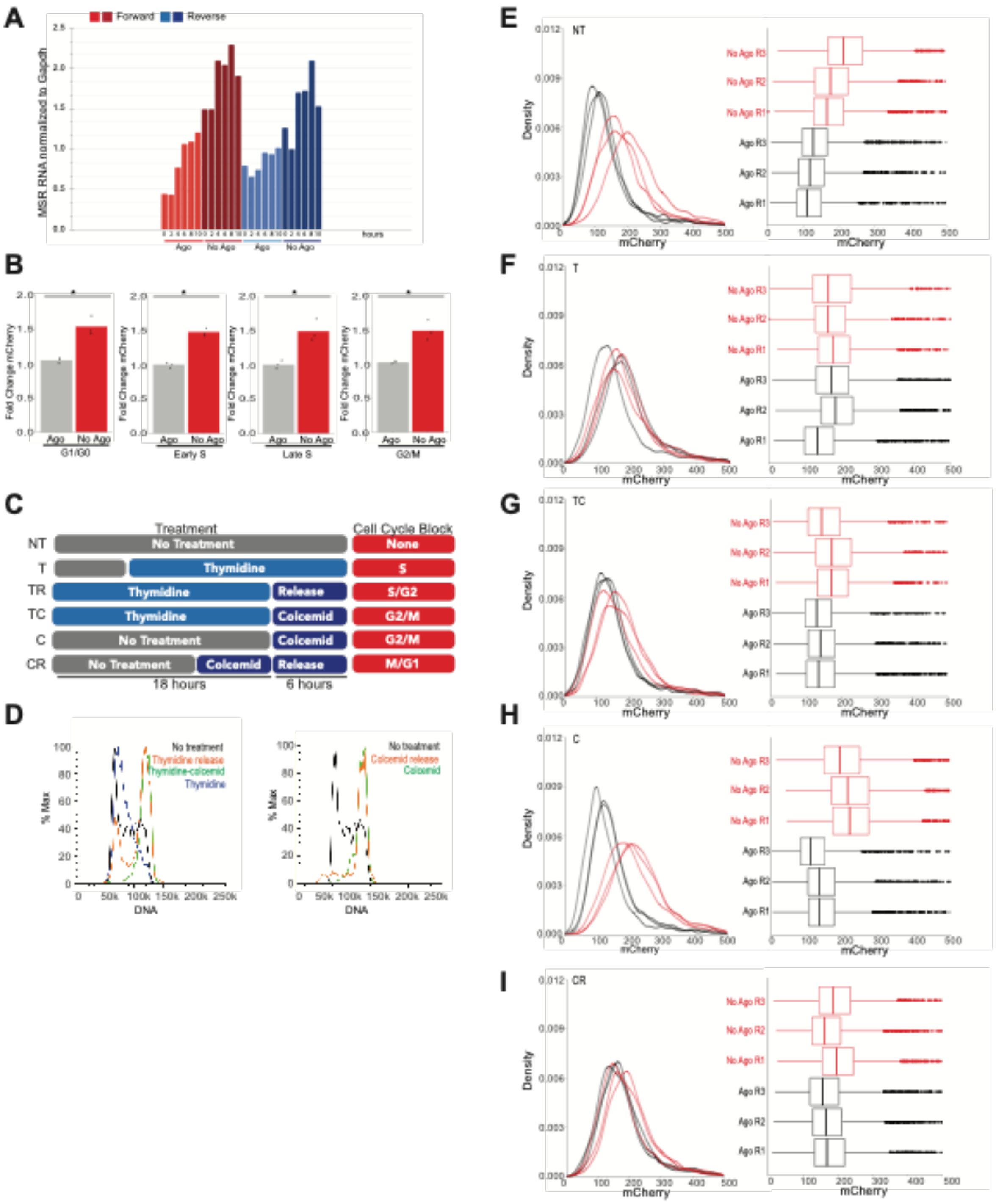

**Supplemental Figure S4. RNAi pericentromeric lncRNA repression occurs after S phase transcription**

(A) Northern quantification of MSR RNA at different time points after release from S phase synchronization. These were performed in cells treated for Ago (Dox) or No Ago (No Dox) expression.

(B) Barplots for the average mCherry expression in MSR2 reporter cells treated for Ago2 (Dox) or No Ago (No Dox) expression and gated based on indicated cell cycle stages. Expression normalized to Ago2-expressing replicate 1 treatment. (* $p$ value $< 0.05$; $p$ value determined by two-sided t-test of biological triplicates)

(C) Diagram of cell cycle treatments used to assay for cell cycle stage regulation.

(D) Histograms confirming enrichment and release from cell cycle treatments.

(E-I) Quantification of mCherry in cells treated for Ago2 (Dox) or No Ago (No Dox) expression and indicated cell cycle treatment performed in biological triplicate.

Sandoval_Figure_S5

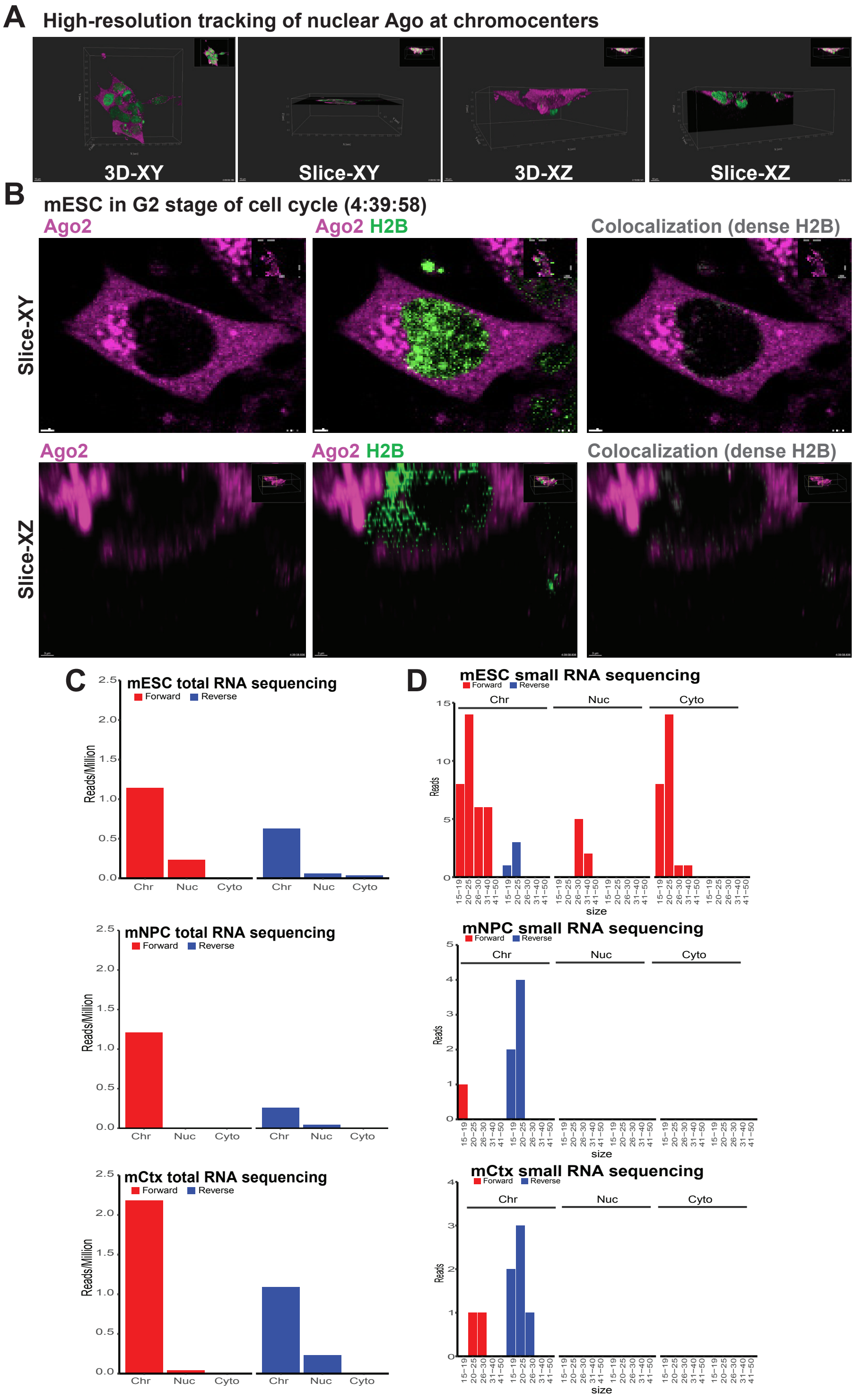

**Supplemental Figure S5. Chromatin-associated pericentromeric lncRNAs are regulated by nuclear Ago**

(A) Images of cellular volume and XY plane (left), and cellular volume and XZ plane (right).

(B) Colocalization of Ago2 with dense H2B-GFP signal in G2 phase of the cell cycle. This is the same dense H2B-GFP region shown in Fig. 5B, but 5 minutes later. See Supplemental Movie 1.

(C) Barplot of mESC fractionated long total RNAseq reads mapping to the MSR. Forward transcripts are in red and reverse in blue. The mESC plot has the same data as Fig. 5D but plotted here on the same scale as mNPC and mCtx for reference.

(D) Barplot of mESC small RNA reads mapping to the MSR in fractionated data. Small RNAs are grouped by size range and plotted separately for forward (red) and reverse (blue) mapping reads. The mESC plot has the same data as Fig. 5E but plotted here on the same scale as mNPC and mCtx for reference.

Sandoval_Figure_S6

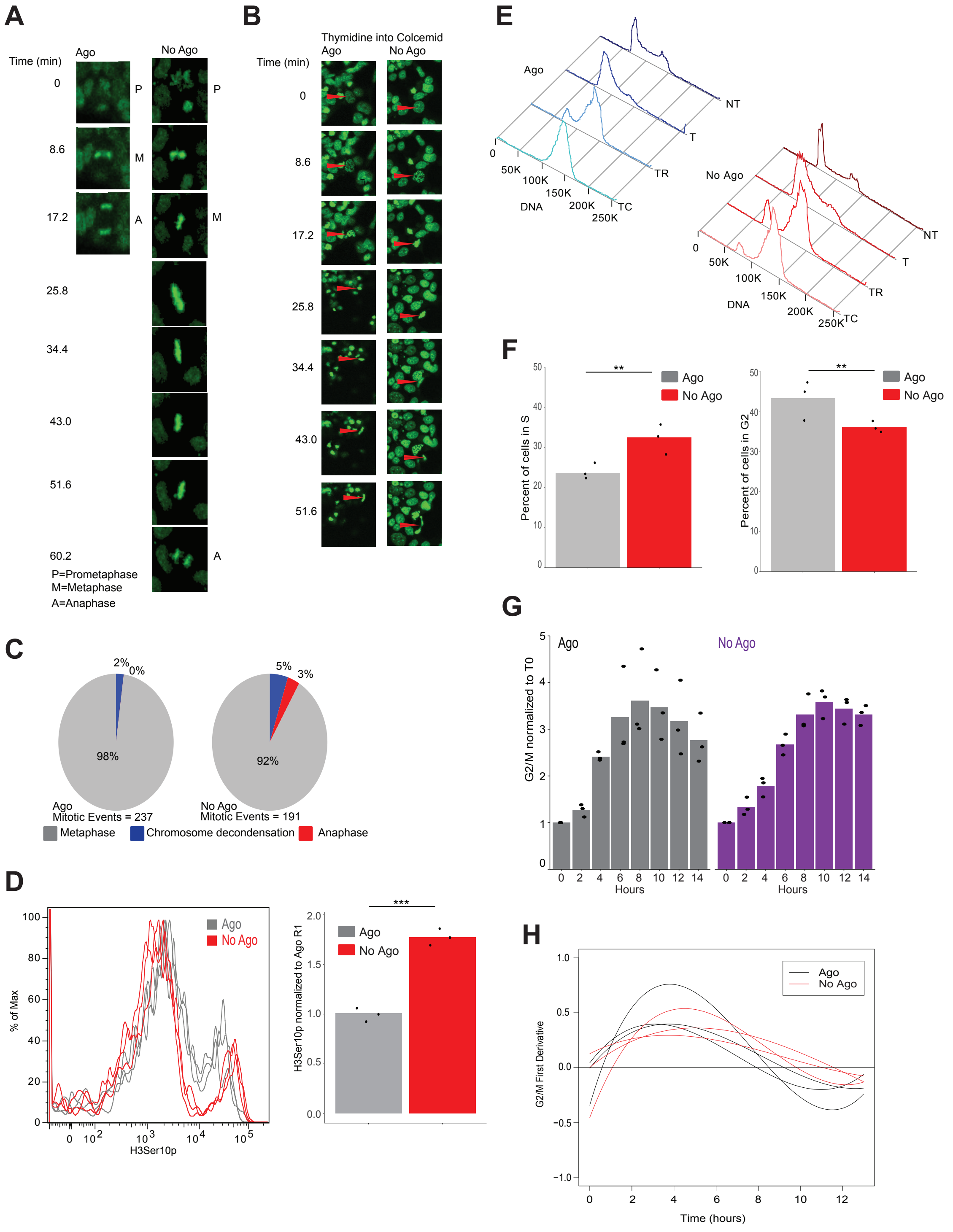

**Supplemental Figure S6. RNAi loss results in mitotic defects that introduce genomic instability**

(A) Example of prolonged metaphase in cells treated for No Ago (No Dox) expression (right) and compared to normal metaphase in cells treated for Ago2 (Dox) expression (left).

(B) Example of SAC defect in cells treated for No Ago (No Dox) expression (right) and compared to normal metaphase arrest in cells treated for Ago2 (Dox) expression (left) after S phase synchronization and released into colcemid for metaphase block.

(C) Quantification of mitotic events using H2B-GFP live-cell imaging for SAC defect after S phase synchronization and released into colcemid for metaphase block.

(D) Quantification of H3Ser10p levels in cells treated for Ago2 (Dox) or No Ago (No Dox) expression. Left: Histogram of H3Ser10p signal after S phase synchronization and released into colcemid for metaphase block. Right: Barplot of average signal normalized to Ago-expressing replicate 1. (***: $p$ value $< 0.001$; p value determined by two-sided t-test of biological triplicates)

(E) DNA profiles for cells treated for Ago2 (Dox) or No Ago (No Dox) expression. NT=no treatment; T=thymidine block; TR=thymidine release; TC=thymidine-colcemid release

(F) Barplots for the average percent of cells in S phase (left) and G2 (right). Cells treated for No Ago (No Dox) or Ago (Dox) expression and thymidine synchronization and release into colcemid. Expression normalized to Ago-expressing replicate 1. (**: $p$ value $< 0.01$; p value determined by two-sided t-test of biological triplicates)

(G) Barplots of the fold increase in G2/M cells compared to amount at release from thymidine (T0). G2/M phase was measured every 2 hours for 14 hours after thymidine release in cells treated for Ago2 (Dox) or No Ago (No Dox) expression. Values are the averages of three experiments each with biological triplicates. Left: Proportion of Ago2-expressing cells (gray). Right: Proportion of Ago-deficient cells (purple). A significant difference between Ago and No No Ago cells is measured at 4 hours using a two-sided t-test.

(H) First derivative of best fit curves for the proportion of G2/M phase cells measured every 2 hours for 14 hours after thymidine release. Ago2-expressing (Dox) cells are represented with black lines and Ago-deficient (No Dox) cells are represented with red lines.

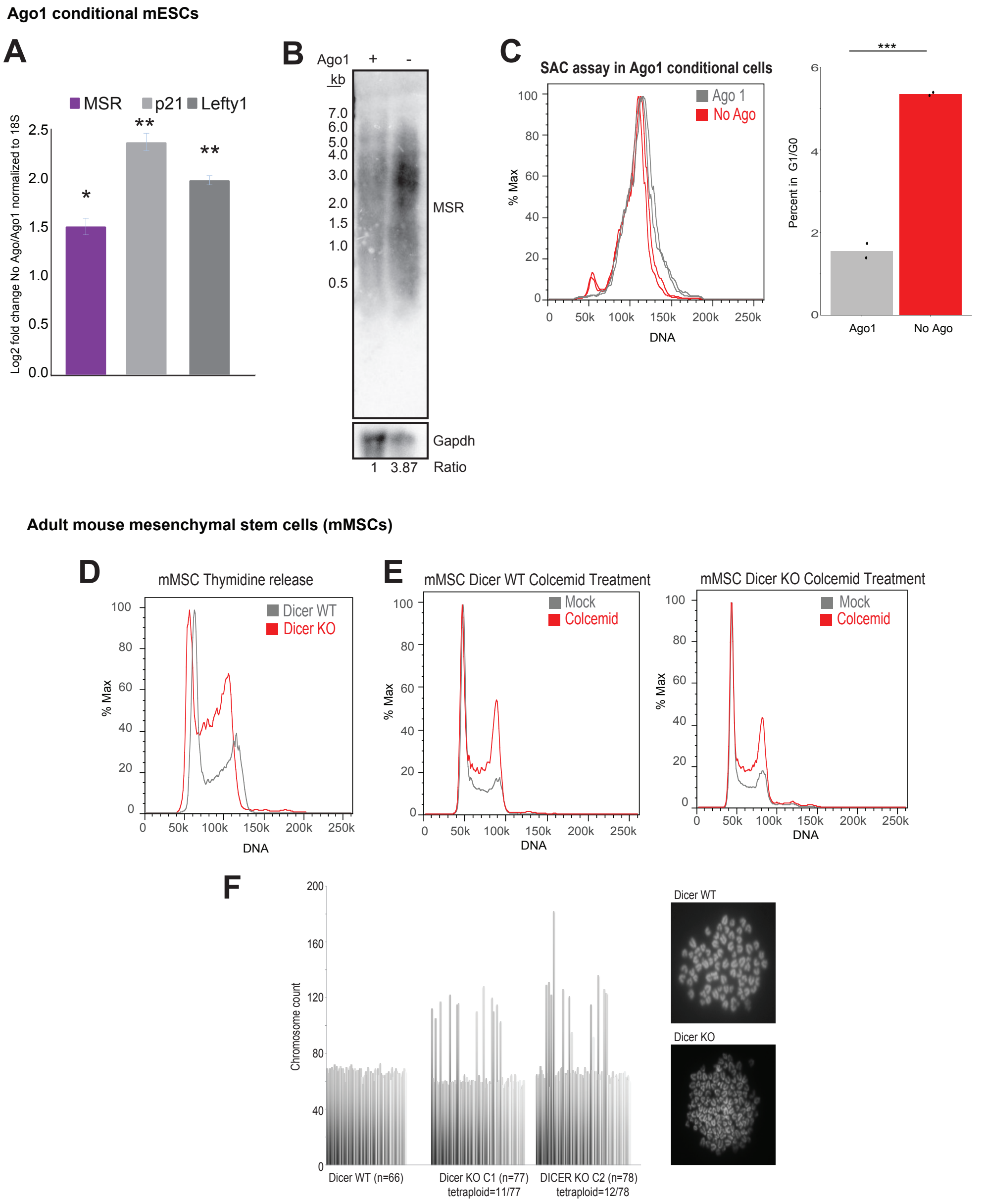
Ago1 conditional mESCs
A
MSR
p21
Lefty1
Log2 fold change No Ago/Ago1 normalized to 18S
*
**
**
B
Ago1 + -
kb
7.0
6.0
5.0
4.0
3.0
2.0
1.5
1.0
0.5
MSR
Gapdh
1 3.87 Ratio
C
SAC assay in Ago1 conditional cells
Ago 1
No Ago
% Max
DNA
***
Percent in G1/G0
Ago1
No Ago
Adult mouse mesenchymal stem cells (mMSCs)
D
mMSC Thymidine release
Dicer WT
Dicer KO
% Max
DNA
E
mMSC Dicer WT Colcemid Treatment
Mock
Colcemid
mMSC Dicer KO Colcemid Treatment
Mock
Colcemid
% Max
DNA
F
Chromosome count
Dicer WT (n=66)
Dicer KO C1 (n=77)
tetraploid=11/77
DICER KO C2 (n=78)
tetraploid=12/78
Dicer WT
Dicer KO

**Supplemental Figure S7. Mitotic defects in other RNAi mutants**
(A) RT-qPCR in cells treated for Ago1 (Dox) or No Ago (No Dox) expression. Quantification of MSR and two miRNA targets (p21 and Lefty1) included. (*: p value < 0.05; **: p value < 0.01; p value determined by two-sided t-test of biological triplicates)

(B) Northern blot of total RNA from cells treated for Ago1 (Dox) or No Ago (No Dox) expression. Top: Random hexamer probe for MSRs; Bottom: Random hexamer probe for Gapdh. Ratio of MSR to Gapdh indicated below.

(C) Left: Histogram of DNA content in cells treated for Ago1 (Dox) or No Ago (No Dox) expression after S phase synchronization and release into colcemid media for SAC assay. Right: Barplot for the average percent of cells in G1/G0. Percent normalized to Ago1 expressing biological replicate 1. (***: p value < 0.001; p value determined by two-sided t-test of biological duplicates)

(D) Delayed progression into G1 shown in histogram of DNA content for mMSCs that are Dicer WT (gray) or Dicer KO (red) after release from thymidine synchronization.

(E) Defective accumulation in metaphase shown in histogram of DNA content for mMSCs that are Dicer WT (left) or Dicer KO (right) with colcemid or mock treated.

(F) Quantification of chromosome numbers in metaphase spreads in mMSC with indicated Dicer genotype.  Right: Example of polyploidy mMSCs in Dicer KO and increased chromosome count in Dicer WT. These viable mMSCs with abnormal chromosome numbers are likely the consequence of altered p53 activity caused by T-antigen transformation used for immortalization.

Table 1. Oligonucleotide Table

| Internal ID | Name | Sequence | Assay | Notes |
|---|---|---|---|---|
| 1H7 | MajorSR3-4_F | aacatccacttgacgacttgaaaaatg | qPCR | Endogenous Major Satellite F |
| 1H8 | MajorSR2_R | ttcctcgccatatttcacgtcctaa | qPCR | Endogenous Major Satellite R |
| 5B7 | 9xMajSat_MSR2_F | TGGCGAGGAAAACTGAAAAAGGT | qPCR | Exogenous Major Satellite F |
| 5B8 | IRES_R | GCTCGTCAAGAAGACAGGGC | qPCR | Exogenous Major Satellite R |
| 4B7 | mGAPDHRT_F | GTGTTCCTACCCCCAATGTGT | qPCR | GAPDH F |
| 4B8 | mGAPDHRT_R | ATTGTCATACCAGGAAATGAGCTT | qPCR | GAPDH R |
| 5B3 | mmRibosomal_18S_F | ATGGTAGTCGCCGTGCCTAC | qPCR | 18S F |
| 5B4 | mmRibosomal_18S_R | CCGGAATCGAACCCTGATT | qPCR | 18S R |
| 4A6 | p21_F | TCCACAGCGATATCCAGACA | qPCR | p21 F |
| 4A7 | p21_R | GGACATCACCAGGATTGGAC | qPCR | p21 R |
| 4E3 | Lefty1_e3e4_F | TCGTGTCCATCCACGAGAGC | qPCR | Lefty1 F |
| 4E4 | Lefty1_e3e4_R | GGGTGCCTTCAGTCACTGGT | qPCR | Lefty1 R |
| 4I4 | Elf_Prom_F, tRNA-glu | ACGCTAGTCAGGTGGGTCAC | qPCR | tRNA-glu |
| 4I5 | Elf_Prom_R, tRNA-glu | GTCAGCCAGAGCCACTAAGC | qPCR | tRNA-glu |
| 5G4 | TTFH MSR BiT FWD | AGA AAA TTG AAA ATT AYG GAA AAT GAG AAA TAT ATA TTT TAG G | PCR | Endogenous Major Satellite F |
| 5G5 | TTFH MSR BiT REV | TTC CAA ATC CTT CAA TAT ACA TTT CAC ATT TTT CA | PCR | Endogenous Major Satellite R |
| 5G6 | 9X MSR BiT FWD 1 | GGATTTGGAATATGGCGAGAAAATTGAAAATTA | PCR | Exogenous Major Satellite F |
| 5G7 | 9X MSR BiT REV 1 | CCCTCACATTACCAAAAAACGACAATATAATAAAA | PCR | Exogenous Major Satellite R |
| 6E1 | MSR Forward ASOligo | mUmAmCmAmGmUmGGACAmUmUmUmCmUmAmA | RNaseH | |
| 6E2 | MSR Reserve ASOligo | mUmUmAmGmAmAmATGTCmCmAmCmUmGmUmA | RNaseH | |
| 6C6 | MSR Forward Probe | GTA AAG TGT GTA TTT CTC | Northern Blot | |
| 6C7 | MSR Reverse Probe | GAG AAA TAC ACA CTT TAC | Northern Blot | |
| 7F9 | 5PRACE_ForMSRprimer_R | GATTACGCCAAGCTTCAGTTTTCCTCGCCATATTTCACGTCCTAAAGTGTGTA | 5' RACE | |
| 7G1 | 3PRACE_ForMSRprimer_F | GATTACGCCAAGCTTCGTGGAATATGGCAAGAAAACTGAAAATCATGGAAAATGAGAAACATC | 3' RACE | |
| 7H6 | 5PRACE_RevMSR | GATTACGCCAAGCTTTGAAAAATGTGAAATGCACACTGAAGGACCTGGAATATG | 5' RACE | |
| 7H7 | 3PRACE_RevMSR | GATTACGCCAAGCTTCACGTCCTAAAGTGTGTATTTCTCATTTTCCGTG | 3' RACE | |
| 1E8 | MajSat_MSR1_sgRNA_F | CACCGAAATGCACACTGAAGGACC | Cas9 targeting | |
| 1E9 | MajSat_MSR1_sgRNA_R | AAACGGTCCTTCAGTGTGCATTTC | Cas9 targeting | |
| 1F1 | MajSat_MSR2_sgRNA_F | CACCGATTTAGAAATGTCCACTGT | Cas9 targeting | |
| 1F2 | MajSat_MSR2_sgRNA_R | AAACACAGTGGACATTTCTAAATC | Cas9 targeting | |
| 5F1 | mmDicer_Bodak_sgRNA1_F | CACCGGGAGTTTAAACCTACAGGCG | Cas9 targeting | Intron 15 |
| 5F2 | mmDicer_Bodak_sgRNA1_R | AAACCGCCTGTAGGTTTAAACTCCC | Cas9 targeting | Intron 15 |
| 3G6 | sgDicer_ex24_4a_F | caccgTCAATAACATGAAAGAGCTC | Cas9 targeting | Exon 24 |
| 3G7 | sgDicer_ex24_4a_R | AAACGAGCTCTTTCATGTTATTGAC | Cas9 targeting | Exon 24 |

**Supplemental_Movie_S1.** Colocalization of Ago2 with dense H2B-GFP heterochromatin in G2 phase into metaphase of mitosis. H2B-GFP in green, Ago2 in magenta, colocalization of Ago2 with dense H2B-GFP signal in gray. See Fig. 5.

**Supplemental_Movie_S2.** H2B-GFP tracking (shown in yellow) in cells treated for No Ago (No Dox) expression. See Fig. 6.

**Supplemental_Movie_S3.** H2B-GFP tracking (shown in yellow) in cells treated for Ago2 (Dox) expression. See Fig. 6.
